# The challenges of independence: ontogeny of at-sea behaviour in a long-lived seabird

**DOI:** 10.1101/2021.10.23.465439

**Authors:** Karine Delord, Henri Weimerskirch, Christophe Barbraud

## Abstract

The transition to independent foraging represents an important developmental stage in the life cycle of most vertebrate animals. Juveniles differ from adults in various life history traits and tend to survive less well than adults in most long-lived animals. Several hypotheses have been proposed to explain higher mortality including that of inadequate/inferior foraging skills compared to adults, young naïve individuals combining lack of experience and physical immaturity. Thus a change in behaviour, resulting in an improvement of skills acquired from growing experience, is expected to occur during a period of learning through the immaturity phase. Very few studies have investigated the ontogeny of foraging behaviour over long periods of time, particularly in long-lived pelagic seabirds, due to the difficulty of obtaining individual tracking data over several years. We investigated the foraging behaviour, through activity patterns, during the three life stages of the endangered Amsterdam albatross by using miniaturized activity loggers on naïve juveniles, immatures and adults. Naïve juveniles during their first month at sea after leaving their colony exhibited lower foraging effort (greater proportion of time spent sitting on water, longer and more numerous bouts on water, shorter and fewer flying bouts). Patterns of activity parameters in juveniles after independence suggested a progressive change of foraging performances during the first two months after fledging. We found sex differences in activity parameters according to time since departure from the colony and month of the year, consistent with the important sexual dimorphism in the Amsterdam albatross. Regardless of life stage considered, activity parameters exhibited temporal variability reflecting the modulation of foraging behaviour. This variability is discussed in light of both extrinsic (i.e. environmental conditions such as variability in food resources or in wind) and intrinsic (i.e. energetic demands linked to plumage renew during moult) factors.

## Introduction

The transition from parental food dependency to independent foraging represents an important developmental stage in the life cycle of most vertebrate animals (Mushinsky et al. 1982; Margrath and Lill 1985; Martin and Bateson 1985; Marchetti and Price 1989; Langen 1996; Burns et al. 2004) and is increasingly documented in a wide range of taxa (reptiles, birds, and some mammals). A widely accepted hypothesis is inadequate/inferior foraging skills of juveniles compared to adults, young naïve individuals combining lack of experience and physical immaturity (Lack 1954; Daunt et al. 2007). Thus, a change in behaviour, resulting from an improvement of skills acquired from increasing experience is expected to occur during a period of learning through the immaturity phase. Learning often refers to stimulus-response associative learning (‘trial and error’; Ruaux et al. 2020), although other forms of learning (such as social learning or imprinting) are also taken into account when considering the ontogeny of complex behaviours (Heyes 1994; Wynn et al. 2020). Such a learning process has been studied on various taxa from insects to primates (Bruner 1972; Caubet et al. 1992; Dukas 2006; Rapaport and Brown 2008).

Juvenile birds are known to undertake vagrant erratic journeys during the post-fledging period in passerines (Naef-Daenzer and Grüebler 2008; Becker 2014; Evans 2018; Boynton et al. 2020), in raptors (Urios et al. 2010; Krüger et al. 2014; Harel et al. 2016) and in seabirds (Riotte-Lambert and Weimerskirch 2013; Collet et al. 2020). Recent studies highlighted that the flight capacities and foraging behaviour of juveniles differed from those of adults in storks (Rotics et al. 2016), raptors (Harel et al. 2016; Nourani et al. 2020) or seabirds (Ydenberg 1989; Péron and Grémillet 2013; de Grissac et al. 2017; Corbeau et al. 2020). Most flight components were found to improve over time to tend towards those of adults (Riotte-Lambert and Weimerskirch 2013; de Grissac et al. 2017; Corbeau et al. 2020).

However, studies focusing on the foraging behaviour of juveniles remain scarce because of the difficulty to obtain individual tracking data for long periods, especially for long-lived pelagic seabirds with deferred maturity. Moreover, existing studies comparing flight capacities and foraging behaviour between juveniles and adults in such species only collected data during the first few months that juveniles spent at sea. Since juveniles may spend several years at sea before returning to a colony to breed, our understanding of the ontogeny of flight capacities and foraging behaviour remains fragmentary.

The Amsterdam albatross *Diomedea amsterdamensis* is a large and long-lived pelagic seabird with an extended immaturity stage (∼ 9 years, Rivalan et al. 2010). Similarly to a closely related species, the wandering albatross *D. exulans*, their foraging strategy relies on very low flight costs as a result of their dynamic soaring flight, whereby individuals optimize the orientation of their movement with wind direction to maximize the daily distance covered (Pennycuick 1982). During initial post-fledging movements juveniles wander alone over very long distances from their colony. At sea distribution during every stage of the life-cycle of Amsterdam albatross was studied by Thiebot et al. (2014) and de Grissac et al. (2016) who compared flight trajectories (i.e. departure direction or orientation toward specific areas) of juveniles and adults. Both studies concluded on slight differences among stages in distribution due to the extensive area they used. However, foraging behaviour is known to be constrained by intrinsic factors such as sex, age, reproductive status and body size across a wide range of taxa and hence play a key role in shaping activity (King 1974; Alerstam and Lindström 1990; Wearmouth and Sims 2008). To understand the changes in foraging proficiency according to experience (life-history stages), longitudinal studies of individuals spanning critical periods of their lives are thus required. Advances in animal-borne instrumentation enable key component of foraging behaviour such as foraging effort and activity to be recorded over long periods.

In this study, we benefited from a unique dataset of different life stages (juveniles, immatures and adults) and a remarkable duration (up to 28 months for juveniles) to characterise and compare the changes in behaviour at sea when birds leave the colony (for several months: immatures and adults, or years: juveniles before returning to land). We analyse the foraging behaviour, through activity patterns, of naïve juveniles (first years of independence at sea), immatures (individuals that never bred, age 2-10 years) and adults (individuals that bred at least once, age 8-28 years) of Amsterdam albatross (Table 1). By using miniaturized activity loggers (Global Location Sensing; GLS) to infer foraging behaviour (activity) throughout the successive life stages we addressed the following questions: i) do individuals belonging to different life-stages behave differently? ii) are there detectable progressive changes in activity patterns? It is noteworthy that the loggers used do not yet allow to have longitudinal data (maximum 2-3 years of recorded data) and to cover the entire period until an individual is recruited into the population as a breeding adult, i.e. at least 8 years.

**Table 1.** Chronological characteristics of life-cycle stages (adapted from Thiebot et al. 2014) and sample sizes of birds tracked using Global Location Sensing (GLS) of Amsterdam albatross.

| Stage<sup>1</sup> | Definition | Age<sup>1</sup> | Tracking duration | Behaviour | Years of deployment | Deployed (n) | Recovered (n) | Recovery rate (%) | GLS with data (n) |
| --- | --- | --- | --- | --- | --- | --- | --- | --- | --- |
| Juvenile | Following chick fledging in January | 1<sup>st</sup> year | ~2.5 years | Chicks disperse at sea after leaving the colony for the first time | 2011 | 21 | 12 | 57 (*t*+9) | 10 (4 F - 6 M)<sup>2</sup> |
| Immature | After juvenile movements, until first breeding attempt (at an average age of 9 years old) | ~2-10 years | ~1 year | Non-breeding young birds forage at sea and occasionally visit the colony for mating | 2011-2012 | 18 | 17 | 94 | 13 (3 F - 9 M – 1 NK) |
| Adult sabbatical | Between two successive breeding periods (~ 15 January year *t* to the following 15 January year *t*+1) | ~8-28 years | ~1 year | Breeding adults at the end of reproductive cycle and leave the colony to forage at sea | 2006, 2009 | 11 | 11 | 100 | 10 (6 F - 4 M) |

<sup>1</sup> Stage/Age at which the individuals were equipped with loggers in our study; <sup>2</sup> number of females F and males M, or not known NK for each stage

Previous knowledge of the ecology of large albatrosses and Amsterdam albatross described above provides a practical framework for testing predictions about variability in foraging behaviour associated with stage, time elapsed since departure from the colony, seasons and sex which are summarised in Table 2. Given the overlap of spatial distribution between life-stages (not presented here but see Thiebot et al. 2014; de Grissac et al. 2016; Pajot et al. 2021) we predicted that juveniles would compensate for any lack of foraging proficiency by increasing foraging effort and time (i.e. lower time spent on water and longer flying bouts, in other words decreasing time sitting on water and increasing number and duration of flight bouts; Hypothesis (A), Table 2). We also predicted changes in activity of juveniles early in post-fledging followed by more progressive changes. Based on results found on wandering albatross fledglings (Riotte-Lambert and Weimerskirch 2013; Pajot et al. 2021) showing that juveniles reached some adult foraging performances in less than two months, we predicted that changes should be detected in activity parameters early after the juvenile left the colony (within few first months). Overall, juveniles should show contrasted foraging effort (i.e. longer time spent on water, shorter flying effort with fewer and shorter flying bouts) early in post-fledging compared to other life-stages. Due to seasonal changes in food availability individuals will face at sea after leaving the colony and the alleviation of energetic constraints linked to reproduction (for breeding adults) or to alternate foraging trips at sea and period on land for pair bonding and mating display (for immature birds), we predicted that adjustments of activity will occur according to the time spent (i.e. in months elapsed) since the departure from the colony (Hypothesis (B), Table 2). In juveniles, we predicted early and rapid changes during post-fledging and then more progressive changes. The two hypotheses (A & B) are not mutually exclusive. While our main objective was to study post-fledging foraging behaviour activity as described above, we also accounted for other sources of changes in foraging behaviour. These included temporal (i.e. related to the month of the year) changes in activity parameters for all life-stages due (i) to environmental changes occurring throughout the seasons, (ii) to partial moulting which is suspected to occur outside the breeding period and to result in reduced activity for adults and immatures (i.e. more time spent on the water; Weimerskirch et al. 2015, 2020), or (iii) to sex differences in flight performances (Shaffer et al. 2001; Riotte-Lambert and Weimerskirch 2013; Clay et al. 2020).

**Table 2.** Hypotheses and predictions about the factors driving differences in activity (time spent on water, number and duration of flying bouts, number and duration of water bouts) year-round in Amsterdam albatrosses. The two hypotheses are not mutually exclusive.

| Hypothesis | Predictions |  |  |
| --- | --- | --- | --- |
|  | (1) Time spent on water (%) | (2) Flying bouts (number/duration) | (3) Water bouts (number/duration) |
| (A) Age and stage specific | Juveniles: increased foraging time/effort and thus lower time spent on water (1), longer flying bouts (2) and shorter water bouts (3) than other stages |  |  |
| (B) Temporal changes - internal requirements: moult/energetic effects | Adults/immatures: two-periods pattern including one with lowering activity |  |  |
|  | Juveniles: change in foraging skills corresponding to gradual change with less time sitting on water (lower time spent on water (1)), increasing flying bouts duration and number (2) and decreasing water bouts duration and number (3), during the 1<sup>st</sup> month after fledging |  |  |
|  | Adjustment in foraging effort to energetic requirements or moult constraints according to time elapsed since departure: higher time spent on water (1), lower flying bouts duration and number (2) and higher water bouts duration and number (3) during moulting |  |  |

## Materials and methods

### Study species and data loggers

Amsterdam Island (37° 50’ S; 77° 33’ E) is located in the subtropical part of the southern Indian Ocean. The Amsterdam albatross, like other great albatrosses, is a biennial breeder (Roux et al. 1983; Jouventin et al. 1989), with high survival during juvenile, immature and adult phase (Rivalan et al. 2010). The adults that raised a chick successfully do not start a new breeding cycle after chick fledging, but remain at sea for a sabbatical period (∼1 yr; Table 1; Rivalan et al. 2010). However, early failed breeders may start to breed the following year (Rivalan et al. 2010). Immature birds may visit the colony when they are 4−7 yrs old, but generally only start breeding at 9 yrs old (Table 1; Weimerskirch et al. 1997a). Juvenile birds fledge and migrate independently from the adults in January (Table 1). Exact fledging dates were not known for juveniles but were assessed from activity pattern as juvenile birds land on water quickly after leaving the colony (Weimerskirch et al. 2006). Amsterdam albatrosses were monitored annually since 1983 and all individuals were individually marked (numbered stainless steel and plastic engraved colour bands; see Rivalan et al. (2010) for details). Unbanded birds of unknown age (79 individuals since the beginning of the study) and chicks of the year were banded, weighed (body mass ± 50 g using a Pesola® spring balance) and measured (wing length ± 1 mm with a ruler, tarsus length, bill length, and bill depth ± 0.1 mm with calipers).

In Amsterdam Island oceanic area, the southern subtropical front (SSTF) delimits the warmer subtropical from the colder sub-Antarctic waters (Belkin & Gordon 1996). Though the diet and foraging strategy of Amsterdam albatross remains poorly known, it is presumed to have very similar foraging behaviour compared to that of the wandering albatross, although subtle differences can appear (Pajot et al. 2021; see Supplementary for species biological aspects). The wandering albatross is known to forage over extensive distances, detecting prey visually or by olfaction during the day (Nevitt et al. 2008) referred as ‘*foraging-in-flight*’, the lowest energy consuming feeding strategy. However, the strategy tends to change depending on breeding stage (Phalan et al. 2007; Louzao et al. 2014), and could result in more frequent and shorter bouts on the water in a ‘*sit and wait*’ technique.

Thiebot et al. (2014) showed that adult Amsterdam albatrosses during their post-breeding sabbatical period moved widely (31° to 115° E), mostly exhibiting westwards wider-scale migratory movements (*sensu* Weimerskirch et al. 2015a) reaching >4000 km from the colony exploiting continuously warm waters (∼18°C; see Supplementary). The immature birds moved widely in longitude (0° to 135° E), exploiting exclusively warm waters 17°-18° C. Juveniles exhibited very large migratory capacities over the southern Indian Ocean after fledging (15° to 135° E, ∼ 4500 km from the colony), through a large range of latitudinal gradient (27° to 47° S). De Grissac et al. (2016) compared trajectories (i.e. departure direction or orientation toward specific areas) of juveniles and adults and showed that juveniles performed an initial rapid movement taking all individuals away from the vicinity of their native colony, and secondly performed large-scale movements similar to those of adults during the sabbatical period.

GLS are archival light-recording loggers used to study activity of birds over periods lasting up to ∼ 2 years. GLSs record the ambient light level every 10 min, from which local sunrise and sunset hours can be inferred to estimate location every 12 h (Wilson et al. 1992). GLS also recorded saltwater immersion data by testing for saltwater immersion at regular interval, storing the number of samples wet (>0) at the end of each 10 min period. We used saltwater immersion to estimate daily activity budget. Despite the higher mean spatial error of location estimates with these devices (over 100 km; Phillips et al. 2004a), GLS loggers allowed us to track the birds for prolonged periods with minimal disturbance to them. We considered the following stages with respect to the ages when GLS were deployed (see Table 1): juvenile, as a fledgling equipped with a GLS just before leaving the colony for the first time; immature, as a non-breeding young bird that had never bred equipped with a GLS when visiting the colony; adult, as a breeding adult equipped with a GLS during the incubation or brooding period which successfully fledged a chick and thereafter took a sabbatical year. To date, we have retrieved 40 of the 50 GLS loggers deployed in total over 4 years, from which 33 individual tracks were estimated (Table 1). Our original aim was to collect activity data on the three stages over a long period of time (>1 year). These data are available from a total of 10 adults tracked throughout their sabbatical period, 13 immature birds and 10 juvenile birds (up to 3.2 years).

### Data processing

The raw immersion data were values from 0 (no immersion or dry, in flight or sitting on the ground) to 200 (permanently immersed in sea water or wet, indicating the number of 3 s periods during 10 min blocks when the sensor was immersed in saltwater). Loggers recorded the proportion of time in seawater at 10 min intervals, which we summarized as hours in the water per day (hereafter time spent on water; 10 min blocks immersion data > 0). This measure is a reliable proxy of foraging effort linked to foraging behaviour of the species which enters the water principally to forage (Weimerskirch and Guionnet 2002). Additionally, the duration of the bouts spent entirely immersed (10 min blocks immersion data = 200) was calculated daily (hereafter referred as wet bouts duration). Conversely, when birds are not on land, the time spent dry was interpreted as flying (and thus not feeding). The duration of the bouts spent entirely dry (10 min blocks immersion data = 0) was calculated daily (hereafter referred as dry bouts duration). Additionally the numbers of bouts (number of wet bouts-sitting on water-and of dry bouts-flying) were obtained daily. Although the loggers integrated activity within each 10 min block and so did not provide the exact timing of landings and take-offs, Phalan et al. (2007) found for comparative purposes that bouts defined as a continuous sequence of 0 values for flight (dry) and a sequence of values of 1 or greater for wet bouts, were suitable proxies for activity.

To select the data corresponding to periods spent at sea after leaving the breeding site, we used the following criteria on activity to define the departure time from the colony for each stage: 1) juveniles, the first bout spent on seawater (wet bouts duration) > 1h based on Argos Platform Transmitters Terminals (PTT) tracking data (data obtained in a other project and not shown here, Weimerskirch et al. unpublished data); 2) immatures and adults, the last bout spent flying (dry bouts duration) > 12h based on PTT tracking data (Weimerskirch et al. unpublished data). Using these criteria we obtained departure months as follows: 1) the juveniles fledged from the colony from January to March, 2) the immatures left between April and August, and 3) the departures of sabbatical adults were spread over two periods, first between December and February and then from May to July.

### Statistical analyses

#### Variation in activity parameters

The aim was to determine whether distinct foraging behaviours could be detected across the patterns of variation of wet/dry data, and then to appraise how these behaviours varied over time and among individuals. First, to deal with the fact that wet/dry metrics were interrelated (number of wet bouts sitting on water and time spent on water, wet bouts duration and dry bouts duration, wet bouts number and dry bouts number) and to avoid redundancy, we ran principal components analyses (PCA built with the ‘PCA’ function, FactoMineR package Lê et al. 2008) to circumvent collinearity issues. To describe changes in behaviour over time and stages using gradients of activity we ran PCA for i) all stages (PCS; based on activity data collected during the first ten months post-departure) and for ii) juveniles only, as an additional goal was to determine changes in activity patterns during the first two years of life (PCJ; based on activity data collected during the first twenty-nine months post-departure).

Considering all stages, the first three principal components (PCS) explained 94.2% of the total variance. For juveniles, the first three principal components (PCJ) explained 92.2% of the total variance. The detailed results of PCA and the variables retained for each axe are summarised in Table 3.

**Table 3.** Results of principal components analyses (PCA) on six wet/dry metrics on Amsterdam albatross.

| Life-stages | Principal components | Total variance explained (%) | Time spent on water | Dry bouts duration | Dry bouts number | Wet bouts duration | Wet bouts number |
| --- | --- | --- | --- | --- | --- | --- | --- |
| All | First (PC1S) | 41.5 | + (r = 0.97)<sup>1</sup> |  | - (r = -0.79) |  |  |
|  | Second (PC2S) | 32.5 |  |  |  | + (r = 0.79) | - (r = -0.75) |
|  | Third (PC3S) | 20.2 |  | + (r = 0.74) | - (r = -0.44) |  |  |
| Juveniles | First (PC1J) | 42.3 | + (r = 0.98) |  | - (r = -0.76) |  |  |
|  | Second (PC2J) | 32.2 |  |  |  | + (r = 0.72) | - (r = -0.75) |
|  | Third (PC3J) | 20.7 |  | + (r = 0.48) | - (r = -0.46) | - (r = -0.46) |  |

<sup>1</sup> the symbol used gives the sign of the correlation (+: positive, -: negative); the number in brackets indicates the value of the correlation coefficient r

Second, we used generalized additive mixed models (GAMMs, built with the ‘gam’ function, itsadug and mgcv package Lin and Zhang 1999; Wood 2015) with the values associated with each of the three first axes of the PCA as the dependent variables. We ran separate models testing for variability in activity parameters i) for all stages combined (PCS) and ii) for juveniles (PCJ), based on different duration of datasets (28 months since departure for juveniles and 9 months since departure for immatures and adults; see Supplementary; Table S1). Thus, for i) we considered the lowest number of months elapsed since departure available (9 months since departure). Months elapsed since departure (the duration elapsed since fledging expressed in month, i.e. the first month after fledging and so on), month of the year (i.e. January and so on), sex, and stage (only for i)) were included as fixed effects. The interactions between stage and time were included as fixed effects to test for the prediction that differences should vanish with time passed since fledging. To test for the importance of individual variability in our results we built models with or without random effects. We compared models without random effect, models with random intercepts, and models with random slopes and intercepts to test whether the rate of change of activity parameters as a function of time elapsed since departure varied between individuals (Zuur 2009a). Models included month elapsed since departure as a continuous covariate modelled with non-parametric smoothing functions (Wood 2017). We limited the amount of smoothing (k) with the ‘gam.check’ function following Wood (2017) for each spline to avoid excessive flexibility and model overfitting that would have no ecological meaning. Models including all combinations of explanatory variables and random effects were then tested and ranked using their Akaike Information Criterion (AIC) values and Akaike weights following the Information-Theoretic Approach (Burnham and Anderson 2002). The model with the lowest AIC was considered as the best model. Two models separated by a difference in AIC values of less than 2 were assumed to fit the data similarly. Using AICc to account for small sample sizes did not change model selection.

Although sexes and stages differed for some body size measurements (see details in Supplementary), we could not include body size as an additional explanatory variable in GAMMs testing for factors of variation in activity patterns due to small sample sizes in each sex and stage category (see Table 1).

Spatial and statistical analyses were performed using R (R Core Team 2021). Values are means ± SD.

## Results

The most parsimonious models explaining variations in activity parameters in the Amsterdam albatross included time elapsed since departure from the colony, month of the year, stages and sexes (Tables 4 and 5; Supplementary Figures S1 - S5; Tables S1), whatever the synthetic activity variables considered (PC1S, PC2S and PC3S; Table 4). The interaction between stage and time elapsed was significant for the first synthetic activity variable (PC1S). Selected models also included random effects on intercepts and slopes, indicating inter-individual variability in activity and in the rate of change of activity as a function of time elapsed since departure from the colony.

**Table 4.** Model selection for variation in activity parameters of Amsterdam albatrosses in relation to sex, stage, number of months spent since departure (month elapsed: duration elapsed since fledging expressed in month, i.e. the first month after fledging and so on) and month of the year (i.e. January and so on)

| Models | Fixed effects | Random effects | AIC | ΔAIC |
| --- | --- | --- | --- | --- |
| Proportion of time spent on water (PC1S) |  |  |  |  |
| M<sub>5</sub> | Month elapsed + Month + Stage + Sex + Month elapsed: Stage | Month elapsed: Individual | 26461.62 |  |
| M<sub>4</sub> | Month elapsed + Month + Stage + Sex | Month elapsed: Individual | 27224.79 | -763.17 |
| M<sub>3</sub> | Month elapsed + Month + Stage | Month elapsed: Individual | 27276.18 | -814.56 |
| M<sub>2</sub> | Month elapsed + Month | Month elapsed: Individual | 28322.25 | -1860.63 |
| M<sub>1</sub> | Month elapsed | Month elapsed: Individual | 28591.69 | -2130.07 |
| M<sub>0</sub> | Null model |  | 28874.42 | -2412.80 |
| Bouts spent on water (PC2S) |  |  |  |  |
| M<sub>3</sub> | Month elapsed + Month + Stage | Month elapsed: Individual | 25751.47 |  |
| M<sub>4</sub> | Month elapsed + Month + Stage + Sex | Month elapsed: Individual | 25752.62 | -1.15 |
| M<sub>2</sub> | Month elapsed + Month | Month elapsed: Individual | 25756.37 | -4.90 |
| M<sub>1</sub> | Month elapsed | Month elapsed: Individual | 25803.80 | -52.33 |
| M<sub>5</sub> | Month elapsed + Month + Stage + Month elapsed: Stage | Month elapsed: Individual | 26750.55 | -999.08 |
| M<sub>0</sub> | Null model |  | 26903.12 | -1151.65 |
| Bouts spent dry -flying (PC3S) |  |  |  |  |
| M<sub>4</sub> | Month + Stage + Sex | Month elapsed: Individual | 22427.29 |  |
| M<sub>3</sub> | Month | Month elapsed: Individual | 22509.79 | -8.14 |
| M<sub>2</sub> | Month elapsed | Month elapsed: Individual | 22539.75 | -82.50 |
| M<sub>1</sub> | Null model | Month elapsed: Individual | 22540.25 | -112.96 |
| M<sub>0</sub> | Null model |  | 23042.26 | -614.97 |

Models are ranked according to decreasing statistical support, as indicated by AIC. The first best models are shown

**Table 5.** Values of activity parameters (mean ± sd) recorded using Global Location Sensing (GLS) depending on stage and sex of Amsterdam albatross.

|  | Juvenile<sup>1</sup> |  | Juvenile<sup>2</sup> |  | Immature |  | Adult sabbatical |  |
| --- | --- | --- | --- | --- | --- | --- | --- | --- |
|  | female | male | female | male | female | male | female | male |
| Time spent on water (%) | 55.04 ± 20.46 | 58.18 ± 21.11 | 51.41 ± 19.18 | 52.88 ± 20.39 | 59.25 ± 21.53 | 63.31 ± 21.17 | 64.89 ± 20.90 | 69.98 ± 18.10 |
| Wet bouts (sitting on water) duration (h) | 1.21 ± 1.74 | 1.24 ± 1.76 | 1.16 ± 1.73 | 1.12 ± 1.59 | 1.07 ± 1.31 | 1.48 ± 2.12 | 1.47 ± 1.95 | 1.33 ± 1.96 |
| Dry bouts duration (h) | 1.29 ± 1.37 | 1.21 ± 1.32 | 1.34 ± 1.41 | 1.26 ± 1.40 | 1.32 ± 1.42 | 1.28 ± 1.55 | 1.44 ± 1.56 | 1.31 ± 1.42 |
| Wet bouts (sitting on water) number | 8.71 ± 4.01 | 8.76 ± 4.09 | 8.14 ± 3.85 | 8.48 ± 4.11 | 10.34 ± 4.29 | 8.59 ± 4.24 | 8.96 ± 3.98 | 10.28 ± 5.33 |
| Dry bouts number | 7.06 ± 3.20 | 7.27 ± 3.52 | 7.57 ± 3.21 | 7.85 ± 3.50 | 6.31 ± 3.21 | 5.75 ± 2.99 | 5.01 ± 2.64 | 4.64 ± 2.48 |

<sup>1</sup> calculated during 28 months following departure; <sup>2</sup> calculated during 9 months following departure

In juvenile Amsterdam albatrosses, the most parsimonious models explaining variations in activity included time elapsed since departure from the colony, month of the year for all three activity variables considered (Table 5 and 6; PC1J, PC2J and PC3J), and sex was retained only for two variables (PC2J and PC3J). Selected models also included random effects on intercepts and slopes, indicating inter-individual variability in activity and in the rate of change of activity as a function of time elapsed since departure from the colony (Supplementary Figures S6).

**Table 6.** Model selection for variation activity parameters for juvenile Amsterdam albatrosses in relation to sex, number of months spent since departure (month elapsed: duration elapsed since fledging expressed in month, i.e. the first month after fledging and so on) and month of the year (i.e. January and so on)

| Models | Fixed effects | Random effects | AIC | ΔAIC |
| --- | --- | --- | --- | --- |
| Proportion of time spent on water (PC1J) |  |  |  |  |
| M<sub>2</sub> | Month elapsed + Month | Month elapsed: Individual | 21625.69 |  |
| M<sub>1</sub> | Month elapsed | Month elapsed: Individual | 21864.11 | -238.42 |
| M<sub>0</sub> | Null model |  | 22109.52 | -483.83 |
| Bouts spent on water (PC2J) |  |  |  |  |
| M<sub>3</sub> | Month elapsed + Month + Sex | Month elapsed: Individual | 19999.00 |  |
| M<sub>2</sub> | Month elapsed + Month | Month elapsed: Individual | 20004.65 | -5.65 |
| M<sub>1</sub> | Month elapsed | Month elapsed: Individual | 20072.42 | -73.42 |
| M<sub>0</sub> | Null model |  | 20417.76 | -418.76 |
| Bouts spent dry -flying (PC3J) |  |  |  |  |
| M<sub>3</sub> | Month + Sex | Month elapsed: Individual | 17541.02 |  |
| M<sub>2</sub> | Month elapsed | Month elapsed: Individual | 17549.00 | -7.98 |
| M<sub>1</sub> | Null model | Month elapsed: Individual | 17548.75 | -7.73 |
| M<sub>0</sub> | Null model |  | 17708.47 | -167.45 |

Models are ranked according to decreasing statistical support, as indicated by AIC. The first best models are show

### Changes in activity for all stages

The two synthetic activity variables (PC1S, PC2S) varied significantly with time exhibiting clear nonlinear temporal patterns (Figure 1). These variations were related to the time elapsed since their departure from the colony and showed seasonal changes (indicated by the month of the year; Supplementary Figures S1 - S5; Tables S1 and S2). With increasing time since departure, birds spent lower percentage of time on water and made shorter wet bouts. They spent less percentage of time on water during the period March to July compared to rest of the year (PC1S, Supplementary Figures S1 - S5). They made longer and fewer bouts on water during the period April to November, and shorter flying bouts during the period November to February. Juveniles showed strong temporal changes in activity linked to the time elapsed since departure from the colony in the first two months after fledging (Supplementary, Figure 2). In immatures and adults the temporal pattern appeared reversed compared to juveniles (Supplementary, Figure 2). Compared to adults, immatures and even more so juveniles, spent a lower percentage of time on water (Table 5, Supplementary Figures S1) and made more flying bouts (PC1S; Supplementary Figures S2), made shorter and fewer bouts on water (PC2S; Supplementary Figures S4-S5), and made longer flying bouts (PC3S; Supplementary Figures S2). Males spent a higher percentage of time on water and made fewer flying bouts (PC1S), longer and more numerous bouts on water (PC2S) and shorter flying bouts (PC3S) compared to females.

**Figure 1.**
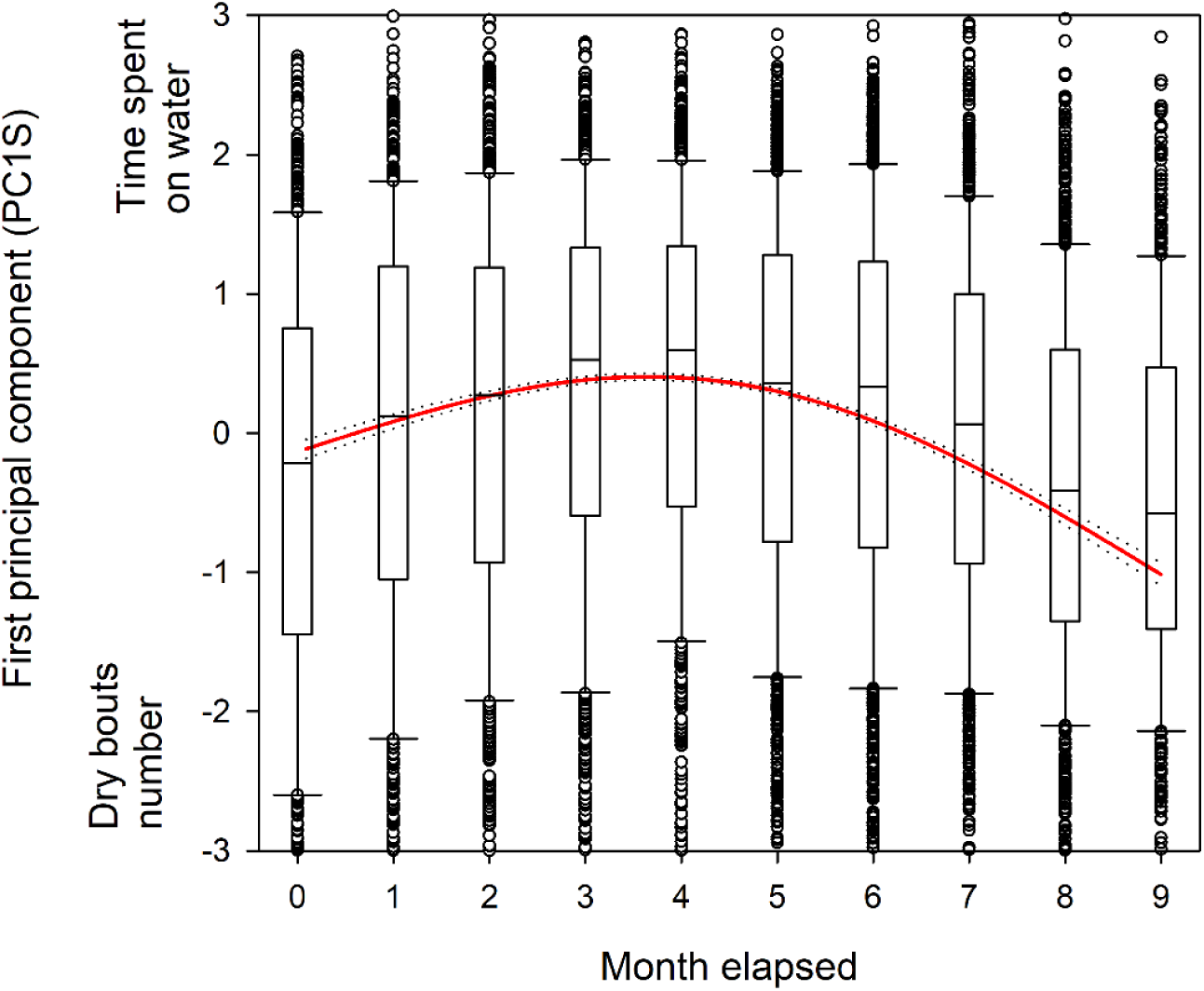
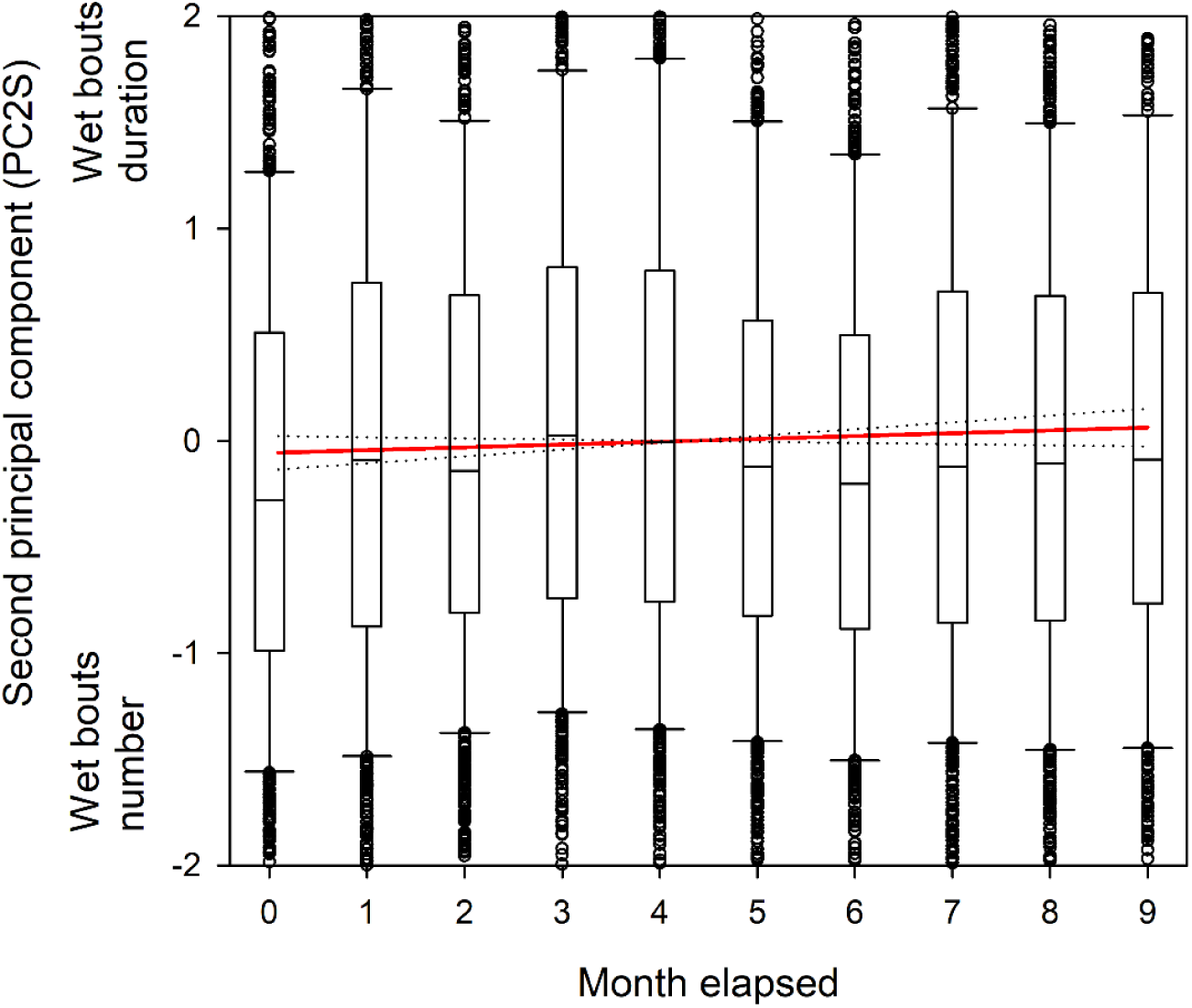
Modeled a) first and b) second axis of principal components analysis of activity parameters for all stages (e.g. adult, immature and juvenile) of Amsterdam albatrosses according to time elapsed (e.g. duration elapsed since departure from the colony expressed in month). Plain line corresponds to estimated smoother from the GAMM model. Dotted lines indicate 95% confidence interval. Boxplot represent raw data. The first axis correlated positively with time spent on water and negatively with dry bouts number and the second axis correlated positively with wet bouts duration and negatively with wet bouts number.

**Figure 2.**
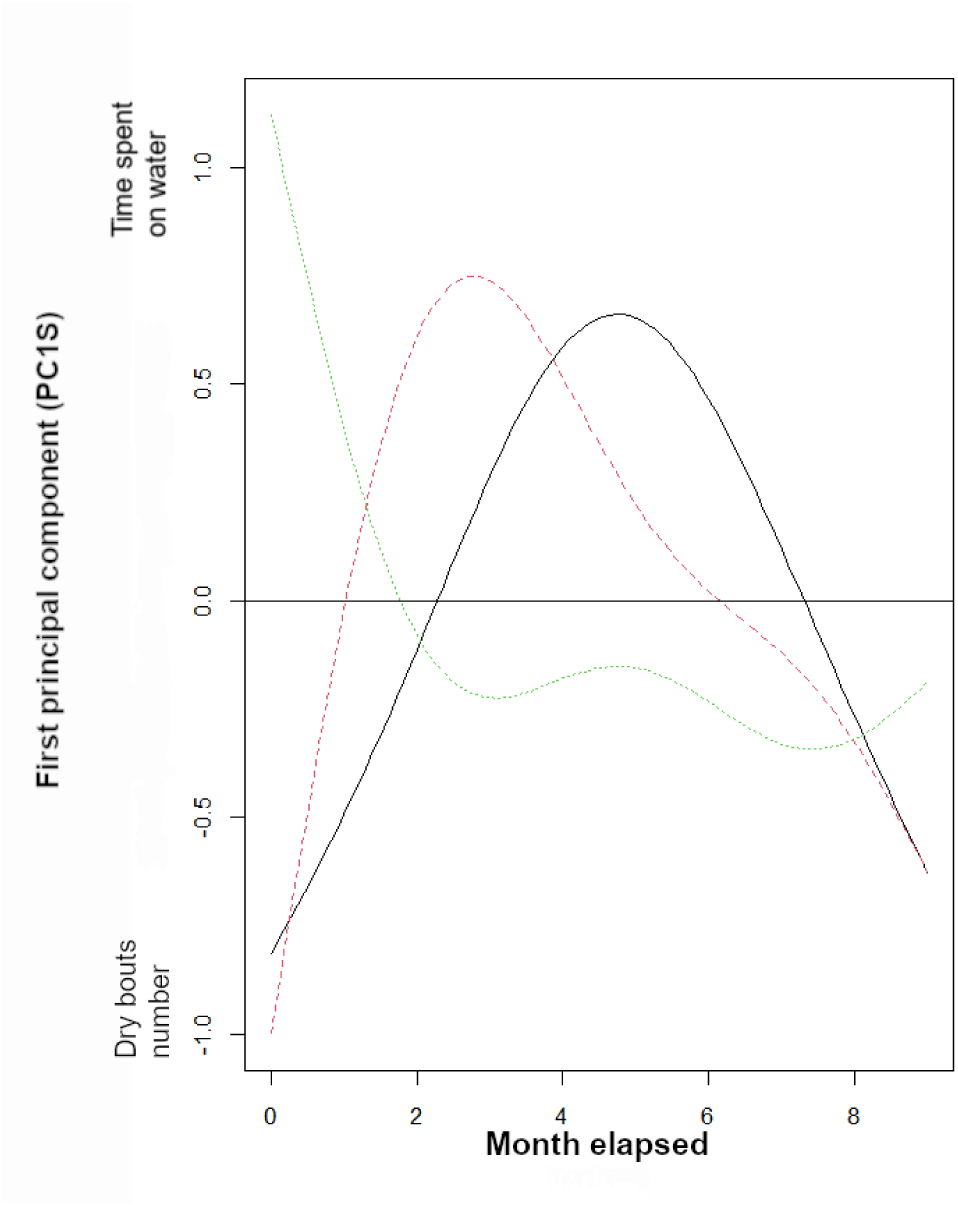
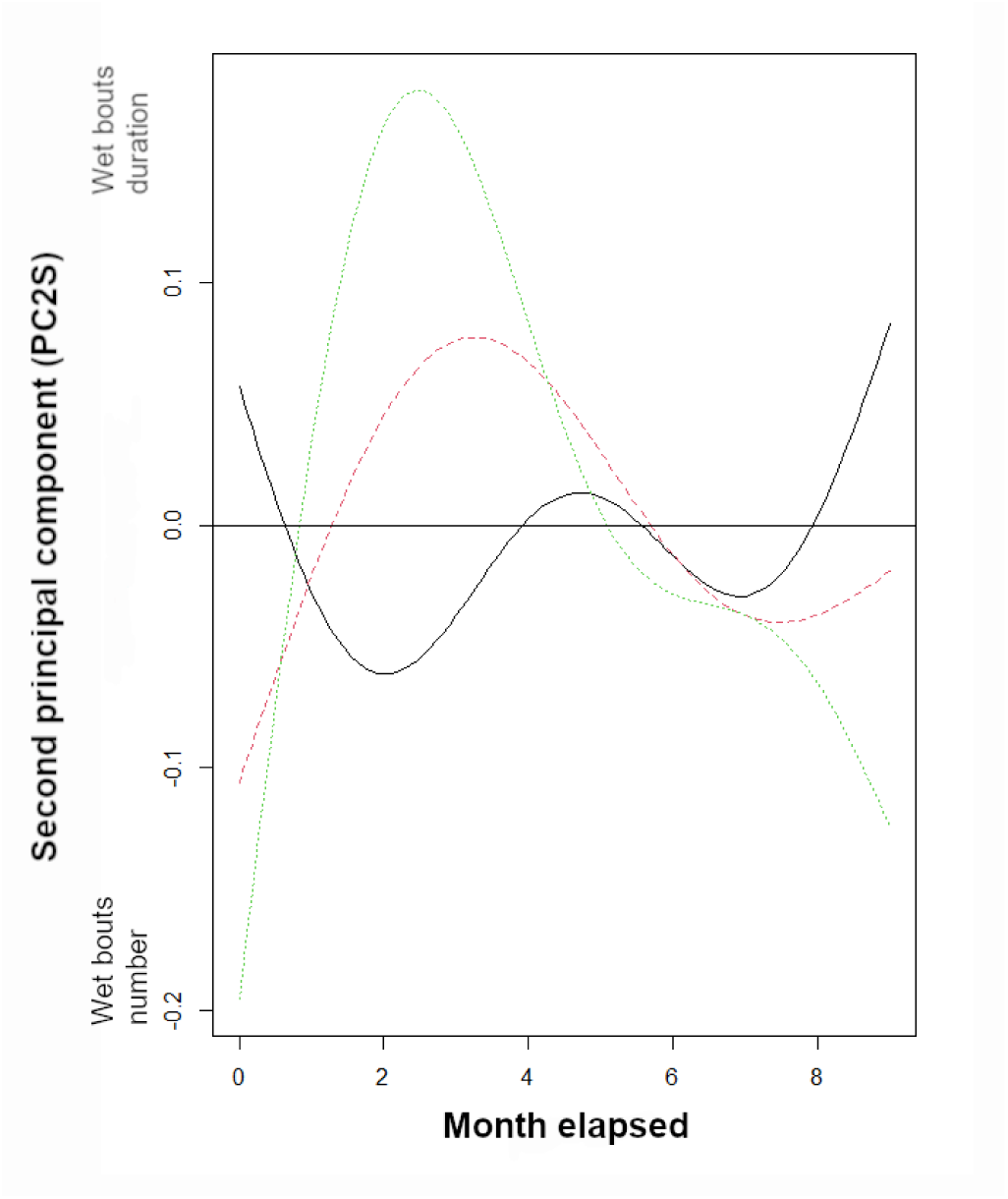
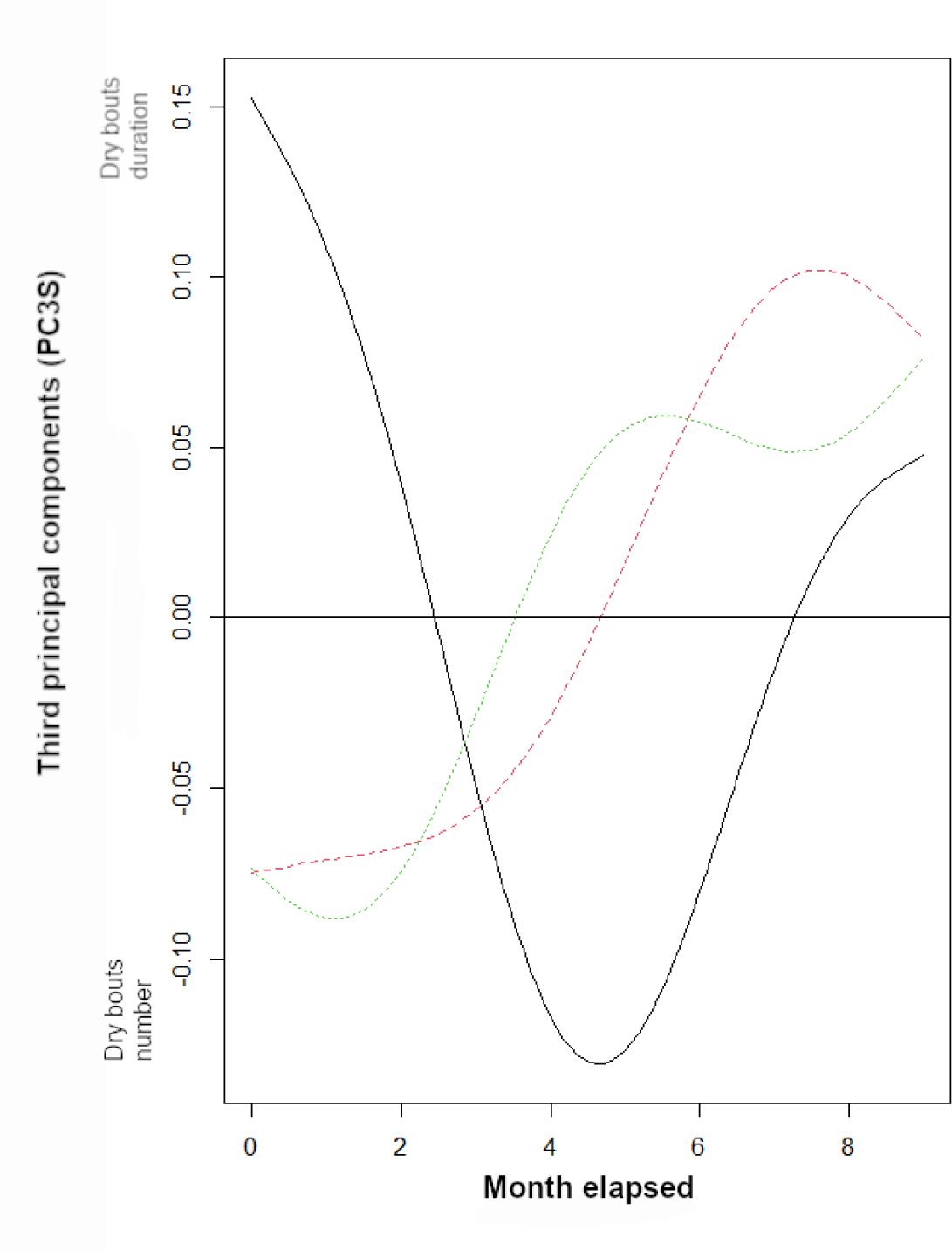
Modeled a) first, b) second and c) third axis of principal components analysis of activity parameters for all stages i.e. adult (plain black line), immature (dashed red line) and juvenile (dotted green line) in Amsterdam albatrosses according to time elapsed (e.g. duration elapsed since departure from the colony expressed in month). Plain line corresponds to estimated smoother from the GAMM model

### Changes in activity of juveniles during the first two years after fledging

PC1J and PC2J varied significantly with time exhibiting clear nonlinear temporal patterns (Figure 3; Supplementary Figures S7 - S11; Tables S1 and S3a, 3b). Juveniles seemed to alternate periods of lower percentage of time spent on water combined with more numerous flying bouts (April) with periods of higher percentage of time on water combined with fewer flying bouts (February, July-October; PC1J, not illustrated). The seasonal change was also observed through longer and fewer bouts spent on water and shorter flying bouts at the end of the year (PC2J: September-December). Juveniles, during the first 28 months after fledging, increased the time spent on water while decreasing the number of flying bouts (Figure 3a).

**Figure 3.**
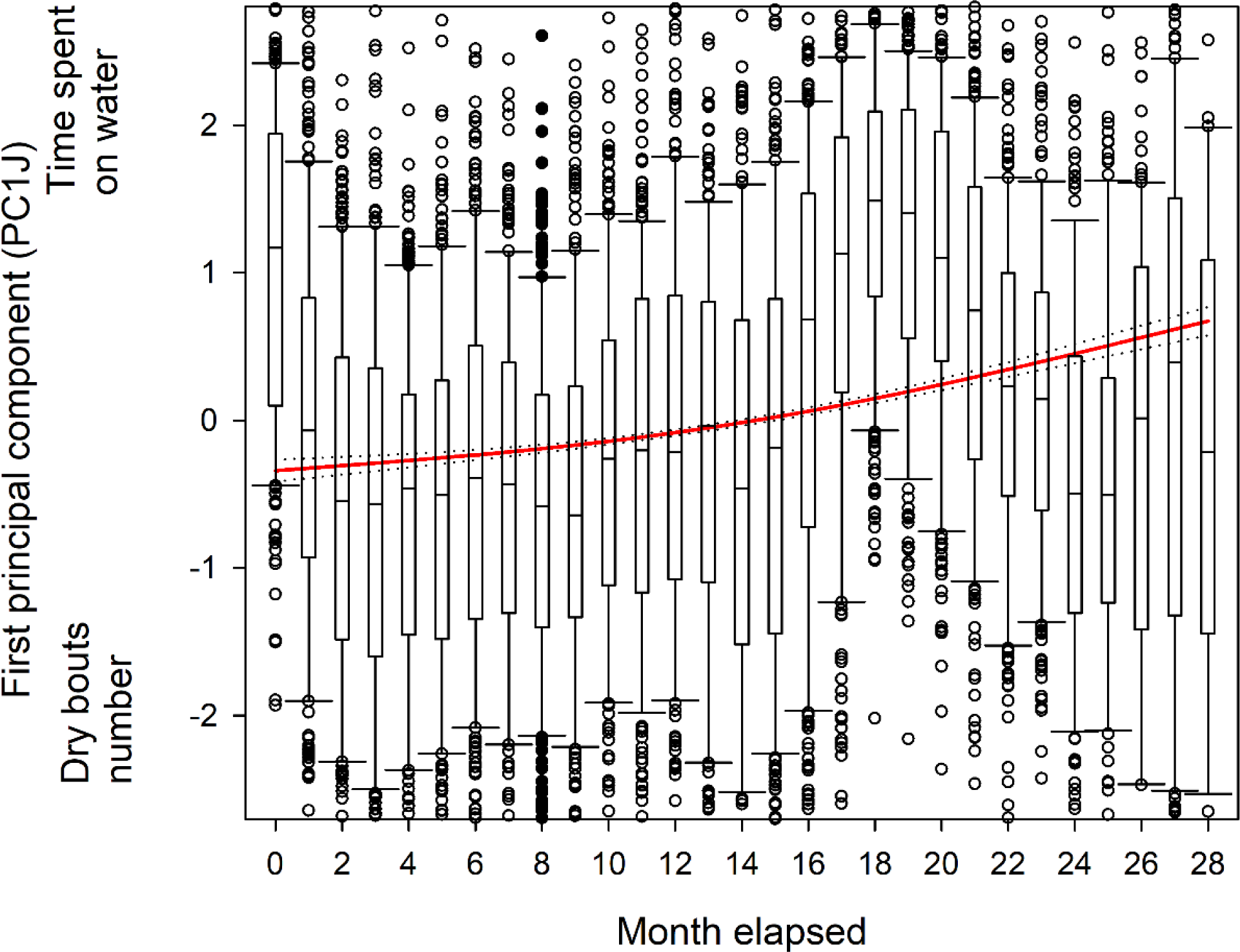
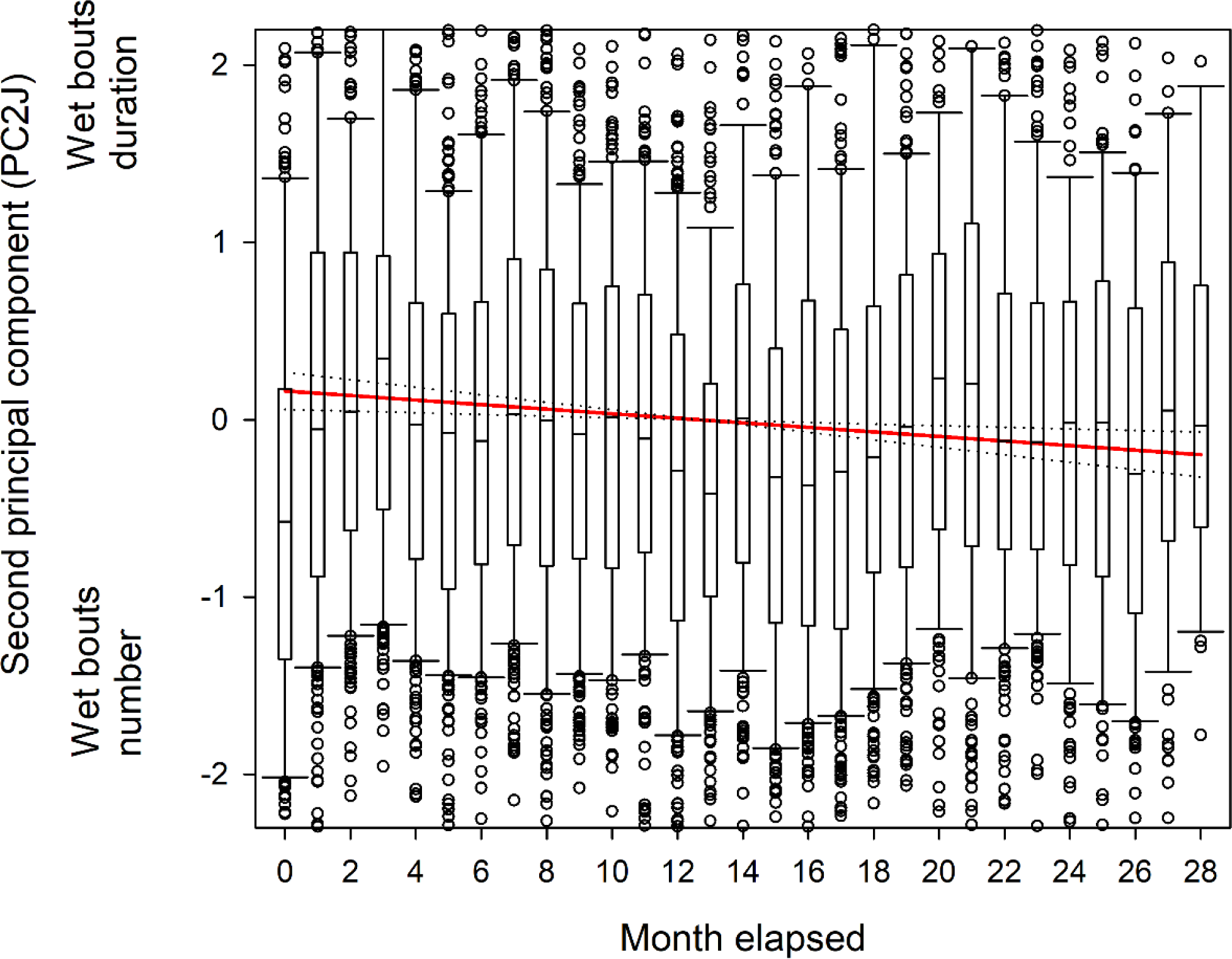
Modeled a) first and b) second axis of principal components analysis of activity parameters of juveniles of Amsterdam albatrosses according to time elapsed (e.g. duration elapsed since departure from the colony expressed in month). Plain line corresponds to estimated smoother from the GAMM model. Dotted lines indicate 95% confidence interval. Boxplot represent raw data.

PC2J and PC3J varied significantly with sex (Supplementary Figures S7 - S10; Tables S5b, 5c), indicating that juvenile males made shorter and more numerous bouts on water (PC2J) and shorter flying bouts (PC3J) compared to females (Supplementary Figures S7 - S10; Tables S5b, 5c).

## Discussion

In this study, we benefited from a unique comprehensive dataset of remarkable duration (up to 28 months) to characterise the post-fledging behaviour of naïve seabirds. Using miniaturized activity loggers (GLS), we showed clear differences and changes in activity characteristics depending on life-stages. By comparing changes in behaviour at sea and foraging parameters of juveniles after their departure at sea with those of immatures and adults in the Amsterdam albatross, we showed that juveniles differed from immatures and adults in their activity values and patterns. Activity also varied according to time and sex. Our study allows us to compare foraging behaviour among life stages in a long-lived endangered seabird species, while also providing new insights into the development of foraging patterns in naïve individuals over a multi-year period.

### Stage specific changes

The birds were found to behave differently according to their stage whatever the activity variables considered, indicating differences in foraging behaviour. Overall, juveniles spent lower percentage of time on water compared to immatures and adults. During the first months following their departure from the colony the proportion of time spent on water by immatures and adults showed a dome-shaped curve peaking three to five months after departure. During the same period of time, the proportion of time spent on water by the juveniles changed strongly, with values dropping off in the first two months and then remaining low and overall lower than in adults and immatures. This might indicate a lower foraging activity in naïve birds. During the same period, the duration and number of water bouts also exhibited progressive change. These patterns suggest an early and gradual change in foraging behaviour so that juveniles progressively could behaved similarly to immatures and adults (reaching similar values in activity covariates). This suggest a progressive behavioural change in movements during the first two months after fledging. It is noteworthy that the multi-monthly bell-shaped pattern observed during the first 10 months after departure in immatures and adults appears to be mirrored in juveniles 15-16 months later (see Figure S11). Despite such possible similarities (% time spent on water), there are still some differences between stages (see Supplementary). Since % time spent on water partly reflects foraging, this seems to indicate that juvenile individuals may have weaker foraging skills during their first two months at sea. Although behavioural changes can often equate to improved performance (e.g. Campioni et al. 2020) this is not always the case. The emergence of juvenile birds as more ‘*adult like*’ in their foraging/flight behavioural patterns is not necessarily a sign of improvement. For example, it could be partly due to individual differences in area use with different environmental conditions encountered (food abundance, wind regimes).

Since all stages of the Amsterdam albatross forage in the same water masses (see Thiebot et al. 2014), differences in foraging behaviour were presumably not due to different oceanographic characteristics as observed in other species (Thiers et al. 2014; Weimerskirch et al. 2014; Frankish et al. 2020b). These differences could be due to a combination of lack of experience, poor knowledge of the environment, use of distinct cues and/or physical immaturity leading to sub-optimal behaviour (Shaffer et al. 2001; Frankish et al. 2020a, 2022). It is likely that increasing exposure to diverse foraging situations allows juveniles to rapidly accumulate foraging experience and improve various aspects of foraging.

Results suggest that immatures may differ from both adults and juveniles in some aspects of their behaviour. While most of the activity parameters and the temporal patterns showed similarities with adults when considering the time elapsed since departure, they seemed rather comparable to juveniles when considering seasonal aspects (month of the year). Such differences can be explained by several non-exclusive explanations: i) similar management of energy constraints than adults, as post-breeding adults and immatures are less constrained in their central-place foraging strategies (Campioni et al. 2016), ii) comparable capacity to respond to local resource availability in their foraging behaviour than juveniles (Frankish et al. 2022), and iii) incomplete acquisition of more long-term learning of complex movement strategies (Thorup et al. 2003; Votier et al. 2011; Rotics et al. 2016). Disentangling these hypotheses can be achieved by combining higher resolution movement data with longer longitudinal studies covering all three life stages for the same individuals.

What might be designated as ‘*lower performance*’ of juveniles found in our study is consistent with studies on wandering albatrosses and Amsterdam albatrosses (Riotte-Lambert and Weimerskirch 2013; de Grissac et al. 2017; Pajot et al. 2021) during the first weeks at sea. Juvenile albatrosses behaved differently in first month after fledging (i.e. speed and sinuosity of movements) and readily use similar foraging strategies as adults (Frankish et al. 2022). Additional skills (such as detection of prey at the surface, detection of other foraging seabirds, navigational skills…) need to be acquired during the immature period before the efficiency of these behaviours matches that of adults. This is also typical of other seabird taxa, which show progressive improvement in flight performance with the numbers of days since fledging (Yoda et al. 2004; Mendez et al. 2017; Collet et al. 2020; Corbeau et al. 2020; Frankish et al. 2022). For example juvenile brown boobies *Anous stolidus* improved their flight abilities (Yoda et al. 2004). In contrast, flight capability (flight speed and sinuosity) comparable to that of adults allows juvenile white-chinned petrels *Procellaria aequinoctialis* to rapidly fly over long distances away from the colony (Frankish et al. 2020). The progressive change of movement behaviours (foraging parameters estimated from activity parameters improved with time elapsed) quantified in juvenile Amsterdam albatrosses could be either due to physical development and/or experience gain. Elucidating the mechanisms of the transition to independence in early life stages is however crucial for understanding the causes of higher juvenile mortality in long-lived species (Fay et al. 2015; Payo-Payo et al. 2016).

### Temporal changes and sex differences in activity

The temporal variability of activity was found whatever the life-stage considered. Part of the activity changes observed following the departure of juvenile Amsterdam albatrosses may illustrate the swift change in travel and movement behaviour, reflecting a more ‘*adult-like*’ behaviour, not indicating necessarily an improvement of flight performances and of the ability to cope with changing (i.e. increasing wind speed) wind conditions (Sergio et al. 2014), a key parameter for soaring seabirds such as albatrosses. Both extrinsic (i.e. environmental conditions) and intrinsic (i.e. energetic demands linked to plumage moult) factors could be involved in the modulation of foraging behaviour, which can be reflected in the temporal variability.

Moult is an intrinsically costly process requiring time, energy and nutrients (Langston and Rohwer 1996; Ellis and Gabrielsen 2002), and the annual replacement of flight feathers is crucial to ensure efficiency in both flight and thermoregulation (Murphy 1996; Peery et al. 2008; Gutowsky et al. 2014). Stage-specific and sex-specific differences in moult extent occur in wandering albatross, suggesting important constraints (Weimerskirch 1991; see Supplementary). Adult birds during the non-breeding season appear to spend much more time on the water during winter, suggesting that partial moult may occur at this time (Weimerskirch et al. 2015b, 2020). Interestingly, immature individuals appear to have a similar peak in time spent on the water in spring, suggesting different timing of moult.

Consistently, we found that males flew for longer periods (dry bouts duration) compared to females. When considering all stages, males spent a higher percentage of time on water compared to females. Males in all stages did more bouts on water and juvenile males shorter wet bouts, compared to females. Contrary to the wandering albatross (Weimerskirch et al. 2014), male and female Amsterdam albatrosses forage in similar oceanic water masses and encounter comparable wind conditions (Jaeger et al. 2013; Thiebot et al. 2014). Therefore, it is unlikely that sex differences in activity parameters were caused by differences in foraging habitats. Sex-specific behavioural differences are common in sexually dimorphic seabirds as reviewed in Wearmouth and Sims (2008), where the smaller sex usually undertakes longer trips. Sexual size dimorphism can result in differences in aerial agility, foraging area and behaviour, and provisioning rate and preferred prey (Gonzales-Solis et al. 2000; Phillips et al. 2004b, 2011; Weimerskirch et al. 2009; Austin et al. 2019; Barbraud et al. 2021). It has also been suggested that size matters probably because the smaller and lighter sex has a higher foraging and flight efficiency (Shaffer et al. 2001; Clay et al. 2020), suggesting that lighter and lower wing loaded female wandering albatrosses, compared to males, are probably better able to exploit subtropical and tropical waters where winds are lighter. Following this, it can be hypothesized that female Amsterdam albatrosses have a greater advantage in foraging in the subtropical environment than males. Sex differences in the acquisition of foraging performance during the first months after fledging yet remain to be explored.

### Individual variability in activity

There was inter-individual variability in almost all activity parameters whatever the stage considered. In juveniles, models indicated inter-individual variability in activity and in the rate of change of activity as a function of time elapsed since departure from the colony. Since the intercept terms in the models were significant, it seems as though individual variability (i.e., specialization on different foraging strategies) was a contributor to observed variability. However, the rate of change of intra-individual variation for some foraging strategies (percentage of time on water-number of flying bouts axis) oscillated during the juvenile period with a seemingly remarkable synchrony (see Fig S7). This suggests that changes in foraging behaviours occurred at the individual level during the juvenile period without stabilizing, at least during the first two years after fledging. This individual variability suggests development of specialized individual foraging behaviours (Harel et al. 2016; Rotics et al. 2016, 2021; Phillips et al. 2017). Nonetheless, given the small sample sizes these results should be interpreted with caution.

## Conclusion

Very few studies have investigated the ontogeny of foraging behaviour over such a long period of time, particularly in long-lived pelagic seabirds, due to the difficulty of obtaining individual tracking data over several years. We investigated the foraging behaviour, through activity patterns, during the three life stages of the endangered Amsterdam albatross by using miniaturized activity loggers. Naïve juveniles during their first month at sea after leaving their colony exhibited lower foraging activity (greater proportion of time spent sitting on water, longer and more numerous bouts on water, and shorter and fewer flying bouts). Patterns of activity parameters in juveniles after independence suggested a progressive change of foraging performances during the first two months after fledging. Regardless of life stage considered, activity parameters exhibited temporal variability reflecting the modulation of foraging behaviour presumably linked to both extrinsic (i.e. environmental conditions such as variability in food resources or in wind) and intrinsic (i.e. energetic demands linked to plumage renew during moult) factors. Sex differences in activity parameters according to time since departure from the colony and season were consistent with the sexual dimorphism in the Amsterdam albatross. Overall, an expected change in behaviour, resulting from the experience gained, may reflect an improvement in skills occurring during a period of learning through the immaturity phase, which would still need to be confirmed by directly assessing foraging performance. Results from our study suggest that the lower foraging activity of juvenile Amsterdam albatrosses may partly explain their lower survival compared to adults. However, despite foraging metrics of juveniles investigated here reached values observed for adults quite rapidly, immature survival remains lower than adult survival during several years (Rivalan et al. 2010). Therefore, more detailed information such as foraging performance realized (the amount or quality of preys obtained by juveniles) or energetic expenditure are needed to better understand age-specific differences in life history traits such as survival and the dynamics of the species.

## Supporting information

data-script

## Ethics

All work was carried out was approved by the French Polar Institute (IPEV) ethics committee.

## Acknowledgements

This study was made possible thanks to all the fieldworkers involved in the monitoring program on Amsterdam albatross, namely Jean-Baptiste Thiebot, Jérémy Demay, Rémi Bigonneau, Romain Bazire, Hélène Le Berre, Marine Quintin, Marine Devaud, Chloé Tanton, Jérémy Dechartre and Anthony Le Nozahic. We acknowledge Dominique Joubert for the management of the demographic CEBC Seabirds database. We are grateful to Richard Phillips, British Antarctic Survey, Cambridge for providing GLS loggers. We thank Bertille Mohring for statistical advices on principal components analyses. We thank our PCI recommender Blandine Doligez and reviewers, Juliet Lamb and an anonymous reviewer for constructive comments on an earlier version of the manuscript.

## Funding

This monitoring program was supported financially and logistically by the French Polar Institute IPEV (program 109, PI C. Barbraud/H. Weimerskirch), the Zone Atelier Antarctique (CNRS-INEE), Terres Australes et Antarctiques Françaises.

This study is a contribution to the National Plan of Actions for Amsterdam albatross.

## Authors contribution

K.D. and C.B. conceived the study. H.W. secured funding. K.D. prepared and analysed the data. C.B. provided feedback on the analyses. K.D. wrote the first draft and all authors contributed to editing versions of the manuscript.

## Supplementary

### Species biological aspects

Though the diet and foraging strategy of Amsterdam albatross remains poorly known, it is presumed to have very similar foraging behaviour compared to that of the wandering albatross, although subtle differences can appear (Pajot et al. 2021). Like other large albatross species (*Diomedea spp.*), the Amsterdam albatross is likely to prey on large squid, fish and carrion found on the sea surface (Delord et al. 2013, Cherel et al. unpublished data). The wandering albatross is known to forage over extensive distances, detecting prey visually or by olfaction during the day (Nevitt et al. 2008). This strategy referred as ‘*foraging-in-flight*’ is the lowest energy consuming feeding strategy for the wandering albatross (Weimerskirch et al. 1997b). However, this strategy tends to change depending on breeding stage (Phalan et al. 2007; Louzao et al. 2014) leading to a more important utilization of the ‘*sit-and-wait*’ technique and possibly to vary depending on sites suggesting considerable behavioural plasticity (Phalan et al. 2007). This switch in foraging techniques could result in more frequent and shorter bouts on the water in the former technique (compared to ‘*foraging-in-flight*’).

Thiebot, et al. (2014) showed that adult Amsterdam albatrosses during their post-breeding sabbatical period moved widely (31° to 115° E), mostly exhibiting westwards wider-scale migratory movements (*sensu* Weimerskirch et al. 2015a) reaching >4000 km from the colony exploiting continuously warm waters (∼18°C). No clear longitudinal seasonality existed in the movements of adults, nonetheless they tended to move westwards in June/July and eastwards in November. The immature birds moved widely in longitude (0° to 135° E), exploiting exclusively warm waters 17°-18° C. Similarly to adults no clear longitudinal seasonality synchronicity existed in the movements, except that they also tended to move westwards in June and eastwards in November. Juveniles exhibited very large post-fledging movement capacities over the southern Indian Ocean after fledging (15° to 135° E, ∼ 4500 km from the colony), through a large range of latitudinal gradient (27° to 47° S). Juveniles birds tended to move westwards first in March-April and temporarily exhibited synchronous individual movements. De Grissac, et al. (2016) compared trajectories (i.e. departure direction or orientation toward specific areas) of juveniles and adults and showed that juveniles performed an initial rapid movement taking all individuals away from the vicinity of their native colony, and in a second time performed large-scale movements similar to those of adults during the sabbatical period. High individual variability and no clear differences between juveniles and adults patterns were found, except that adults foraged at significantly higher latitudes. De Grissac et al. (2016) concluded in an overlap in distribution between adults and juveniles due to the extensive area they used and their differences in latitudinal distribution compared to other Procellariiformes species.

Moult is an intrinsically costly process requiring time, energy and nutrients (Langston and Rohwer 1996; Ellis and Gabrielsen 2002), and the annual replacement of flight feathers is crucial to ensure efficiency in both flight and thermoregulation (Murphy 1996; Peery et al. 2008; Gutowsky et al. 2014). In large-sized albatrosses like Amsterdam albatross, replacement of primary feathers lasts for more than one breeding season, and the moult of primaries never occurs during the breeding season (Furness 1988; Weimerskirch 1991). Stage-specific and sex-specific differences in moult extent occur in wandering albatross, suggesting important constraints that could compete with breeding (immature birds tend to renew fewer feathers compared to adult breeders), and particularly in females (Weimerskirch 1991). In smaller sized seabirds, a link between moulting pattern and activity parameters was evidenced, resulting in a clear temporal pattern partly explained by moult (Cherel et al. 2016). Recently Gutowsky et al. (2014) suggested that tropical albatrosses (i.e. Laysan *Phoebastria immutabilis* and black-footed *P. nigripes* albatrosses) could compromise flight from active wing moult during the nonbreeding period and induce changes in daily activity budget during a ‘quasi-flightless’ stage. However, there is no such data for southern albatrosses. Furthermore for large sized species (*Diomedea spp.*) the activity data recorded using GLS never suggested it such a compromise. However, adult birds during the non-breeding season appear to spend much more time on the water during winter, suggesting that partial moult may occur at this time, as observed in many other seabird species that have to moult during the non-breeding season and show reduced activity during specific periods that may correspond to moulting (Weimerskirch et al. 2015b, 2020).

### Statistical analyses

#### Variation in activity parameters between stages with time-lag

The visual comparison shown on Figure S11 was statistically tested using generalized additive mixed models (GAMMs, built with the ‘gam’ function, itsadug and mgcv package, (Lin and Zhang 1999; Wood 2015)) with the values associated with the first axe of the PCA as the dependent variable. We ran model testing for variability in activity parameters for all stages combined (PC1Slag; Table S4). We applied time lag as illustrated in Figure S11, the first axe was modelled as a function of months spent since departure from the colony (monthelap.lag) with a delay of 16 months.

#### Variation in body size

Differences between sexes in body size measurements were tested using Student’s t-tests and Wilcoxon rank tests. We tested independently if each measurement (wing length, tarsus length, bill length, bill depth and body mass) varied according to sex and stage (juvenile and adult). The effects were tested using generalised linear models (GLMs) with a Gaussian family and identity link function (Zuur 2009b). Model validation and model selection were performed following (Zuur 2009b). GLMs tested for effect of sex and stage and T-tests tested the differences of body size measurements between males and females. Although sexes and stages differed for some body size measurements, we could not include body size as an additional explanatory variable in GAMMs testing for factors of variation in activity patterns due to small sample sizes in each sex and stage category.

Male Amsterdam albatrosses were larger than females, particularly for tarsus length and bill length and bill depth whatever the stage (juvenile or adult; Tables S5). In juveniles, males were ∼13% heavier than females, while the difference was not significant in adults (Table S5). The most sexually dimorphic phenotypic traits were body mass, bill depth and tarsus length in juveniles while in adults they were body mass, tarsus length and bill length.

---

**Table S1.** Selected models testing for the effects of sex, stage, number of months spent since departure (monthelap: duration elapsed since fledging expressed in month, i.e. the first month after fledging and so on) and month of the year (i.e. January and so on) on activity parameters of Amsterdam albatrosses.

|  | Model # | Study variable<sup>1</sup> | Model structure | Sample size |
| --- | --- | --- | --- | --- |
| All stages | gamm1 | PC1S | ~s(monthelap, by=stage, k = 2) + monthf + stade + sex + s(monthelap, device_code<sup>2</sup>, bs = "re") | 8094 |
| All stages | gamm2 | PC2S | ~ s(monthelap, k = 3) + monthf + stade + s(monthelap, device_code, bs = "re") | 8094 |
| All stages | gamm3 | PC3S | ~monthf+stade+sex+s(monthelap,device_code, bs='re') | 8094 |
| Juveniles | gamm4 | PC1J | ~ s(monthelap,k=2)+monthf+s(monthelap,device_code, bs='re') | 6161 |
| Juveniles | gamm5 | PC2J | ~ s(monthelap, k = 2)+monthf+sex+s(monthelap, device_code, bs = "re") | 6161 |
| Juveniles | gamm6 | PC3J | ~monthf+sex+s(monthelap,device_code, bs='re') | 6161 |

<sup>1</sup> First, second and third principal component issued from principal components analyses considering i) all stages combined (PCS) and ii) only
juveniles (PCJ); <sup>2</sup> Individuals

**Table S2a.**
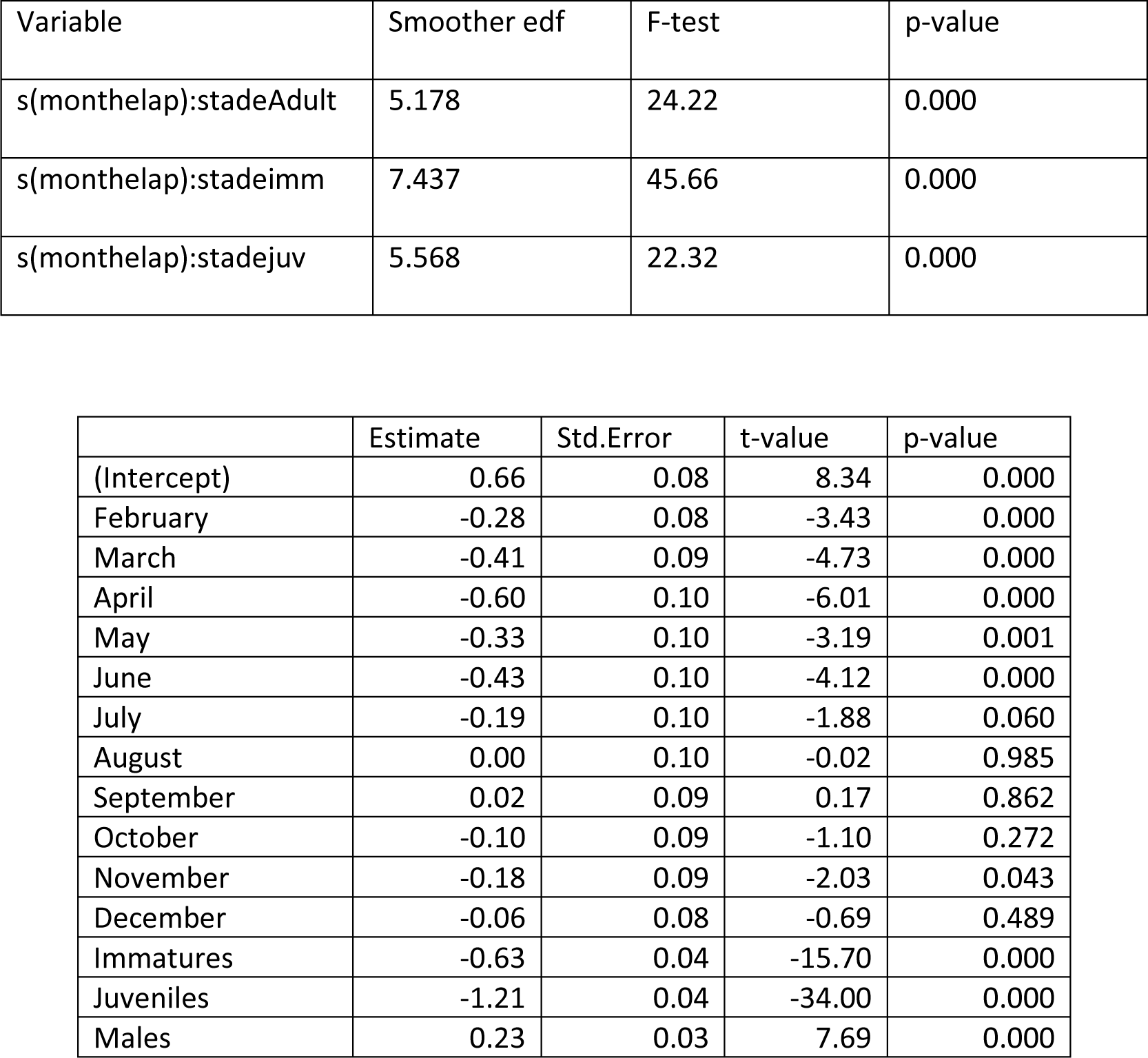
GAMM results for the first principal components (PC1S; gamm1 see Table S2) of Amsterdam albatross modelled as a function of months spent since departure from the colony (monthelap), month of the year, stage and sex. Reference values are January, adults and females.

| Variable | Smoother edf | F-test | p-value |
| --- | --- | --- | --- |
| s(monthelap):stadeAdult | 5.178 | 24.22 | 0.000 |
| s(monthelap):stadeimm | 7.437 | 45.66 | 0.000 |
| s(monthelap):stadejuv | 5.568 | 22.32 | 0.000 |

|  | Estimate | Std.Error | t-value | p-value |
| --- | --- | --- | --- | --- |
| (Intercept) | 0.66 | 0.08 | 8.34 | 0.000 |
| February | -0.28 | 0.08 | -3.43 | 0.000 |
| March | -0.41 | 0.09 | -4.73 | 0.000 |
| April | -0.60 | 0.10 | -6.01 | 0.000 |
| May | -0.33 | 0.10 | -3.19 | 0.001 |
| June | -0.43 | 0.10 | -4.12 | 0.000 |
| July | -0.19 | 0.10 | -1.88 | 0.060 |
| August | 0.00 | 0.10 | -0.02 | 0.985 |
| September | 0.02 | 0.09 | 0.17 | 0.862 |
| October | -0.10 | 0.09 | -1.10 | 0.272 |
| November | -0.18 | 0.09 | -2.03 | 0.043 |
| December | -0.06 | 0.08 | -0.69 | 0.489 |
| Immatures | -0.63 | 0.04 | -15.70 | 0.000 |
| Juveniles | -1.21 | 0.04 | -34.00 | 0.000 |
| Males | 0.23 | 0.03 | 7.69 | 0.000 |

**Table S2b.** GAMM results for the second principal components (PC2S; gamm2 see Table S2) of Amsterdam albatross modelled as a function of months spent since departure from the colony (monthelap), month of the year, stage and sex. Reference values are January, adults and females.

| Variable | Smoother edf | F-test | p-value |
| --- | --- | --- | --- |
| s(monthelap) | 1.001 | 0.504 | 0.478 |
| s(monthelap,device_code) | 27.107 | 39.991 | 0.000 |

|  | Estimate | Std.Error | t-value | p-value |
| --- | --- | --- | --- | --- |
| (Intercept) | -0.15 | 0.10 | -1.53 | 0.126 |
| February | 0.13 | 0.08 | 1.71 | 0.088 |
| March | 0.16 | 0.08 | 2.00 | 0.046 |
| April | 0.42 | 0.08 | 5.03 | 0.000 |
| May | 0.40 | 0.08 | 4.99 | 0.000 |
| June | 0.25 | 0.08 | 3.16 | 0.002 |
| July | 0.23 | 0.08 | 2.92 | 0.004 |
| August | 0.26 | 0.08 | 3.40 | 0.001 |
| September | 0.48 | 0.08 | 6.22 | 0.000 |
| October | 0.35 | 0.08 | 4.57 | 0.000 |
| November | 0.34 | 0.08 | 4.41 | 0.000 |
| December | 0.19 | 0.08 | 2.49 | 0.013 |
| Immatures | -0.12 | 0.08 | -1.57 | 0.116 |
| Juveniles | -0.18 | 0.06 | -2.96 | 0.003 |

**Table S2c.** GAMM results for the third principal components (PC3S; gamm3 see Table S2) of Amsterdam albatross modelled as a function of months spent since departure from the colony (monthelap), month of the year, stage and sex. Reference values are January, adults and females.

| Variable | Smoother edf | F-test | p-value |
| --- | --- | --- | --- |
| s(monthelap,device_code) | 26.52 | 16.58 | 0.000 |

|  | Estimate | Std.Error | t-value | p-value |
| --- | --- | --- | --- | --- |
| (Intercept) | 0.34 | 0.06 | 5.37 | 0.000 |
| February | -0.22 | 0.06 | -3.43 | 0.000 |
| March | -0.07 | 0.06 | -1.08 | 0.279 |
| April | -0.10 | 0.07 | -1.53 | 0.127 |
| May | 0.00 | 0.06 | 0.05 | 0.958 |
| June | 0.05 | 0.06 | 0.87 | 0.385 |
| July | 0.02 | 0.06 | 0.39 | 0.694 |
| August | -0.04 | 0.06 | -0.70 | 0.483 |
| September | -0.06 | 0.06 | -0.93 | 0.355 |
| October | -0.10 | 0.06 | -1.58 | 0.012 |
| November | -0.16 | 0.06 | -2.57 | 0.010 |
| December | -0.23 | 0.06 | -3.70 | 0.000 |
| Immatures | -0.27 | 0.06 | -4.61 | 0.000 |
| Juveniles | -0.45 | 0.05 | -9.12 | 0.000 |
| Males | -0.14 | 0.04 | -3,39 | 0.000 |

**Table S3a.**
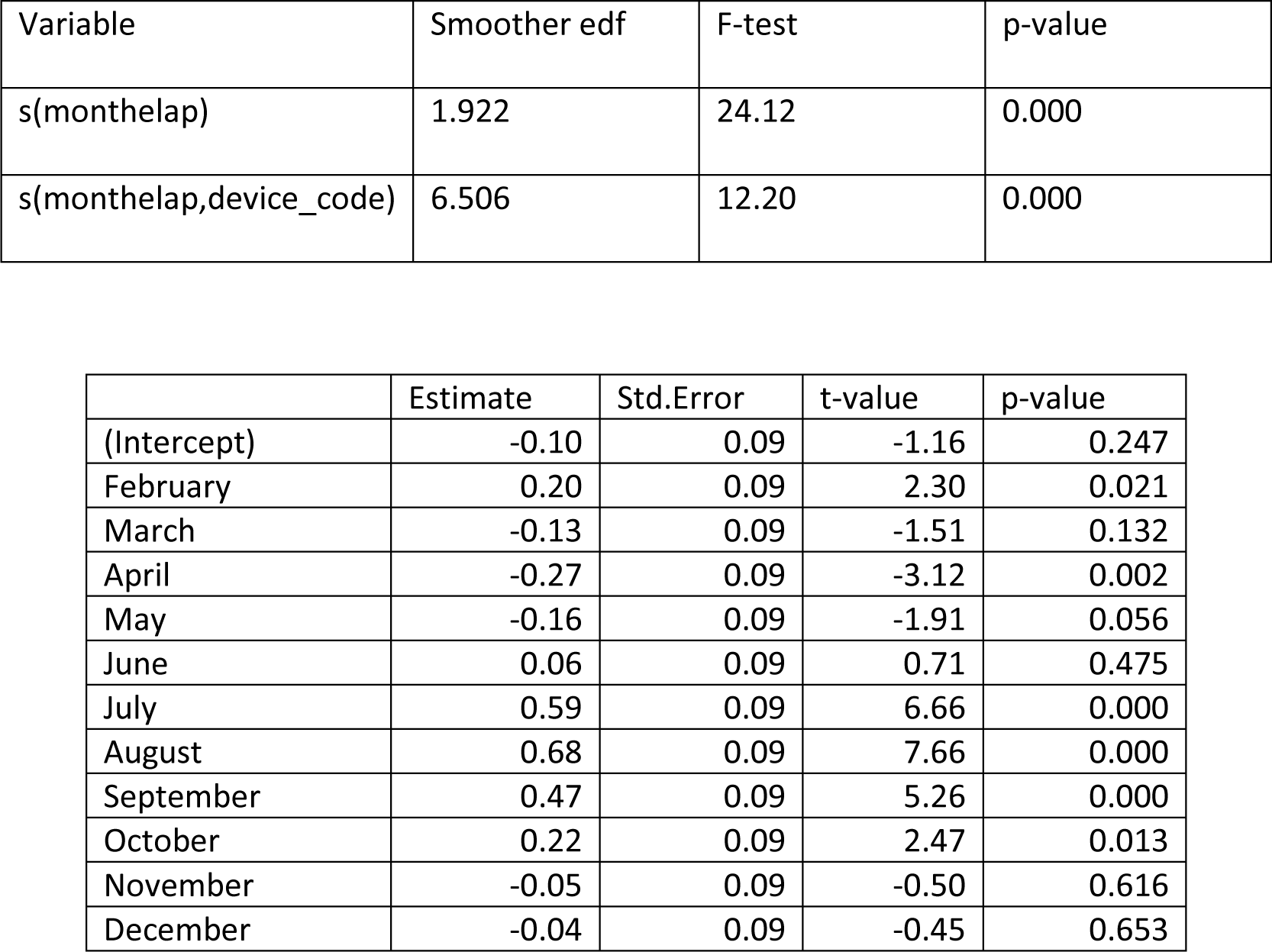
GAMM results for the first principal components (PC1J; gamm4 see Table S2) of juveniles Amsterdam albatross modelled as a function of months spent since departure from the colony (monthelap) and month of the year. Reference value is January.

**Table S3b.**
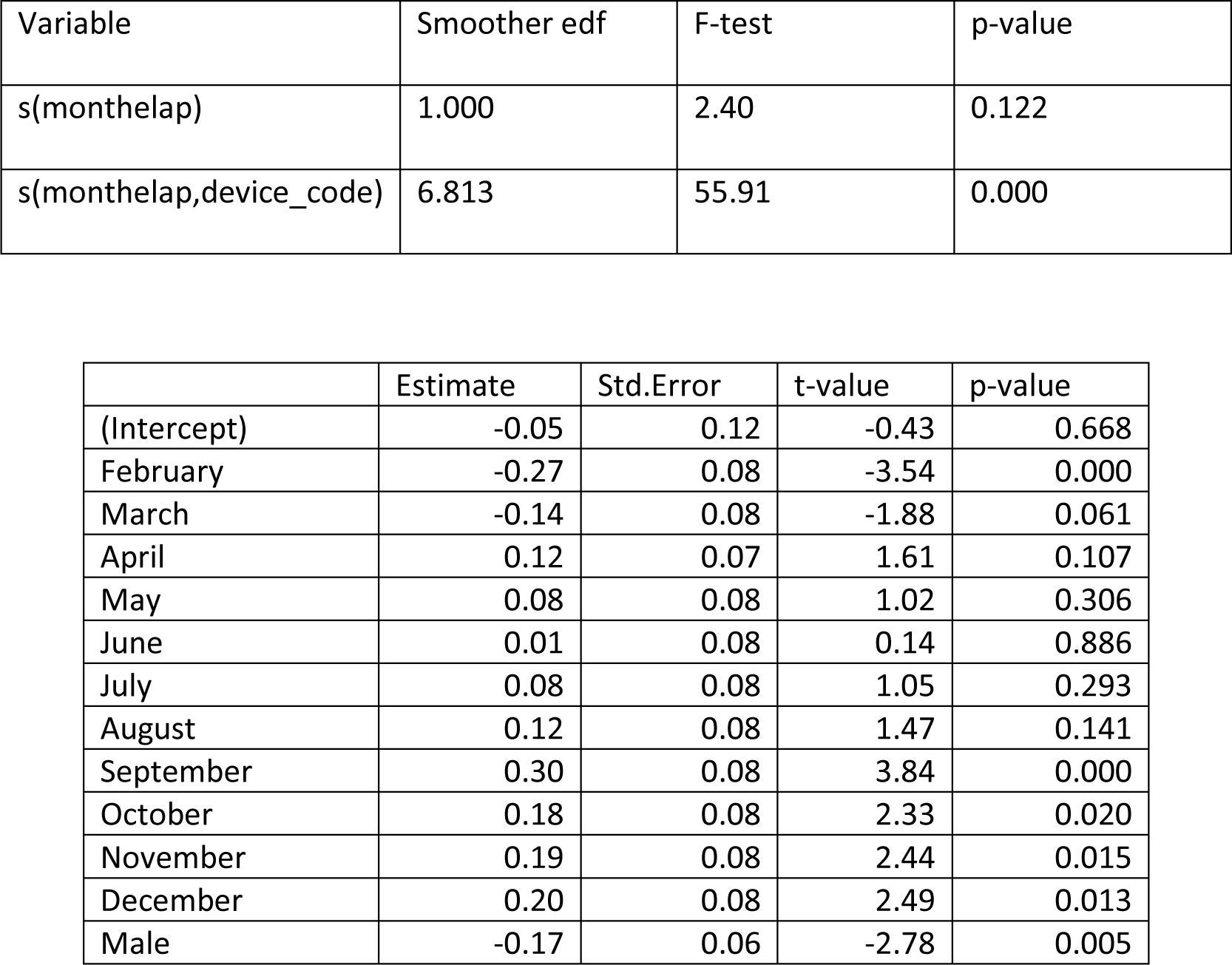
GAMM results for the second principal components (PC2J; gamm5 see Table S2) of juveniles Amsterdam albatross modelled as a function of months spent since departure from the colony (monthelap) and month of the year. Reference value is January.

**Table S3c.**
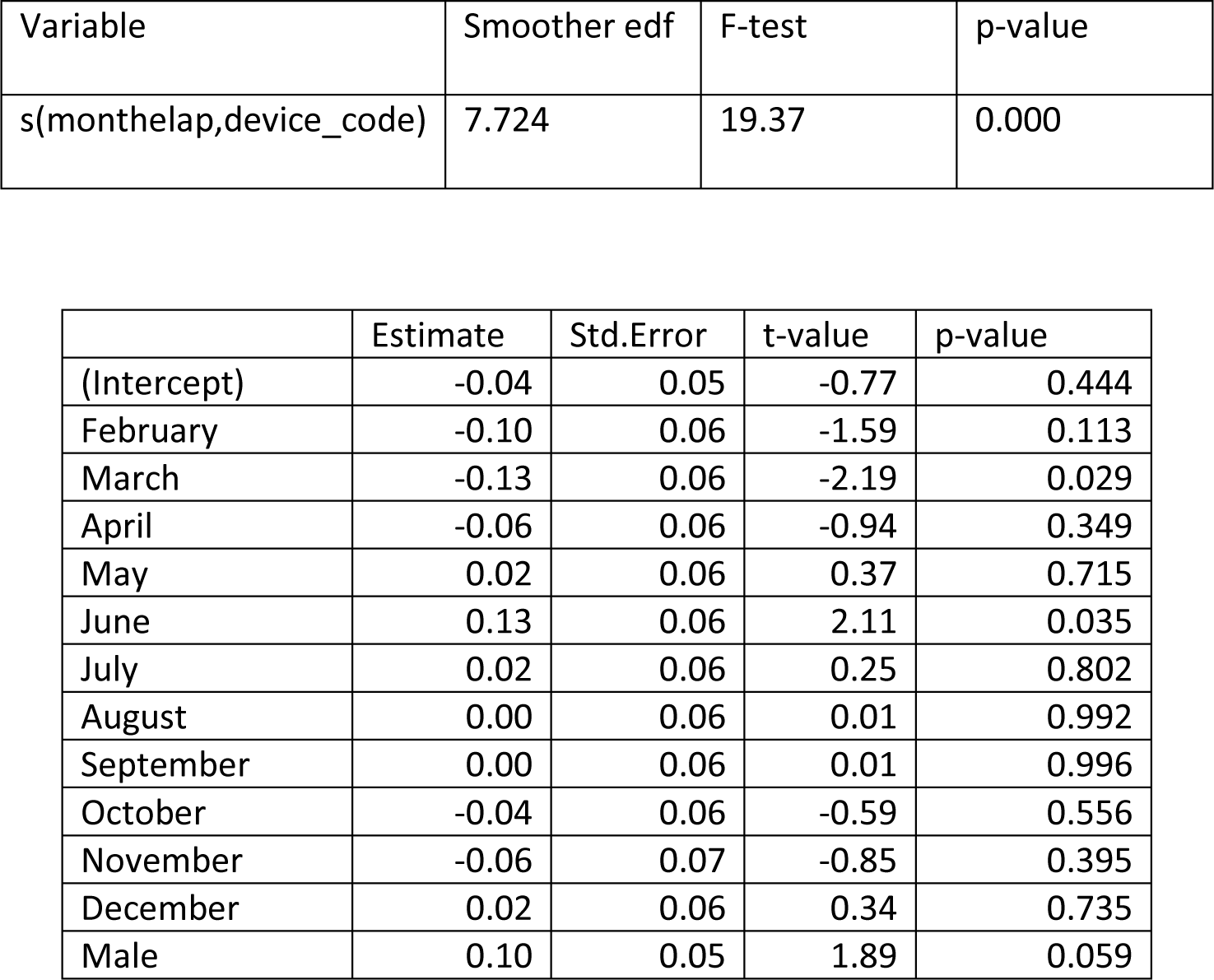
GAMM results for the third principal components (PC3J; gamm5 see Table S2) of juveniles Amsterdam albatross modelled as a function of months spent since departure from the colony (monthelap), month of the year and sex. Reference value are January and females.

| Variable | Smoother edf | F-test | p-value |
| --- | --- | --- | --- |
| s(monthelap,device_code) | 7.724 | 19.37 | 0.000 |

|  | Estimate | Std.Error | t-value | p-value |
| --- | --- | --- | --- | --- |
| (Intercept) | -0.04 | 0.05 | -0.77 | 0.444 |
| February | -0.10 | 0.06 | -1.59 | 0.113 |
| March | -0.13 | 0.06 | -2.19 | 0.029 |
| April | -0.06 | 0.06 | -0.94 | 0.349 |
| May | 0.02 | 0.06 | 0.37 | 0.715 |
| June | 0.13 | 0.06 | 2.11 | 0.035 |
| July | 0.02 | 0.06 | 0.25 | 0.802 |
| August | 0.00 | 0.06 | 0.01 | 0.992 |
| September | 0.00 | 0.06 | 0.01 | 0.996 |
| October | -0.04 | 0.06 | -0.59 | 0.556 |
| November | -0.06 | 0.07 | -0.85 | 0.395 |
| December | 0.02 | 0.06 | 0.34 | 0.735 |
| Male | 0.10 | 0.05 | 1.89 | 0.059 |

**Table S4.** GAMM results for the first principal components (PC1Slag) of Amsterdam albatross modelled as a function of months spent since departure from the colony (monthelap.lag) with a delay of 16 months (see Figure S11), month of the year, stage and sex. Reference values are January, adults and females.

| Variable | Smoother edf | F-test | p-value |
| --- | --- | --- | --- |
| s(monthelap.lag):stadeAdult | 5.001 | 49.37 | 0.000 |
| s(monthelap.lag):stadeimm | 4.810 | 19.39 | 0.000 |
| s(monthelap.lag):stadejuv | 7.643 | 53.53 | 0.000 |

|  | Estimate | Std.Error | t-value | p-value |
| --- | --- | --- | --- | --- |
| (Intercept) | 0.99 | 0.08 | 11.95 | 0.000 |
| February | -0.53 | 0.09 | -5.79 | 0.000 |
| March | -1.08 | 0.09 | -11.74 | 0.000 |
| April | -1.48 | 0.09 | -15.75 | 0.000 |
| May | -1.23 | 0.09 | -13.59 | 0.001 |
| June | -1.03 | 0.09 | -11.82 | 0.000 |
| July | -0.42 | 0.08 | -4.92 | 0.060 |
| August | -0.07 | 0.08 | -0.90 | 0.985 |
| September | 0.02 | 0.08 | 0.28 | 0.862 |
| October | -0.05 | 0.08 | -0.70 | 0.272 |
| November | -0.15 | 0.08 | -1.96 | 0.043 |
| December | 0.05 | 0.08 | 0.67 | 0.489 |
| Immatures | -0.81 | 0.06 | -14.24 | 0.000 |
| Juveniles | -0.74 | 0.05 | -14.71 | 0.000 |
| Males | 0.20 | 0.03 | 6.96 | 0.000 |

**Table S5.**
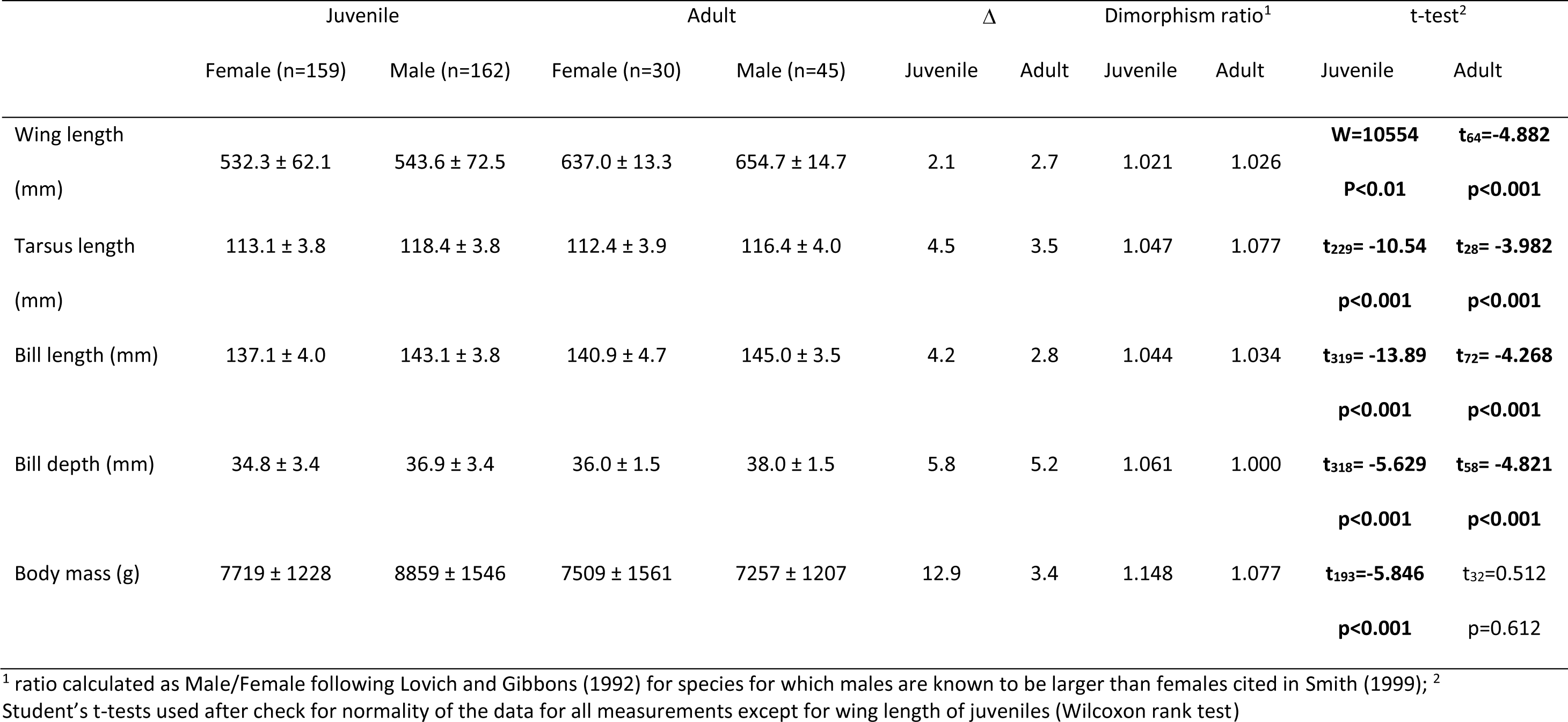
Body measurements of juveniles and adults Amsterdam albatross and percentage of differences between sexes for each measurement. Δ is the difference in %, p values are reported.

## FIGURES

**Figure S1.**
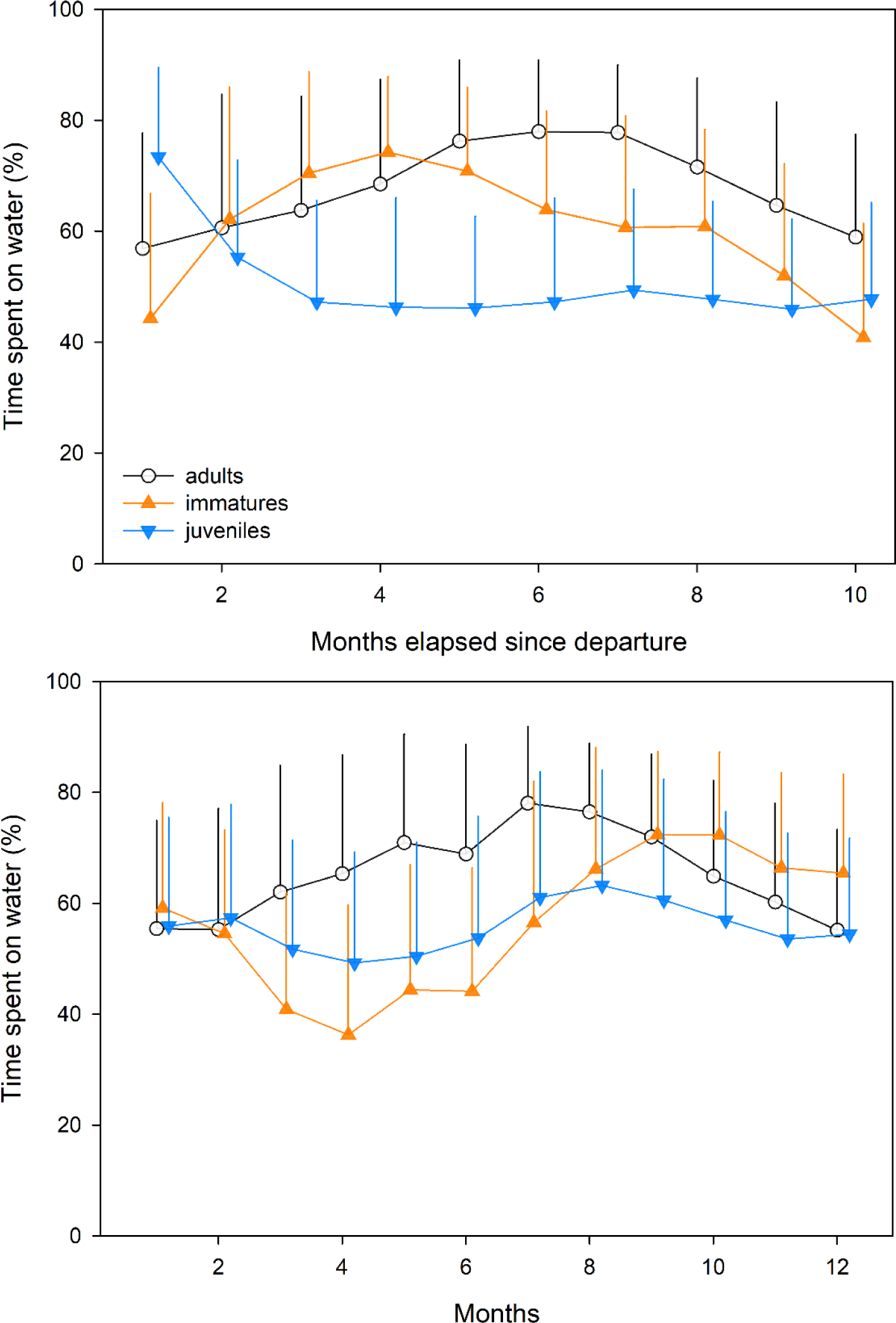
Daily proportions of time spent on water depending on stage (juveniles, immatures and adults) for every month since departure from the colony (upper panel) and for each month of the year (lower panel). Error bars represent ± 1 sd

**Figure S2.**
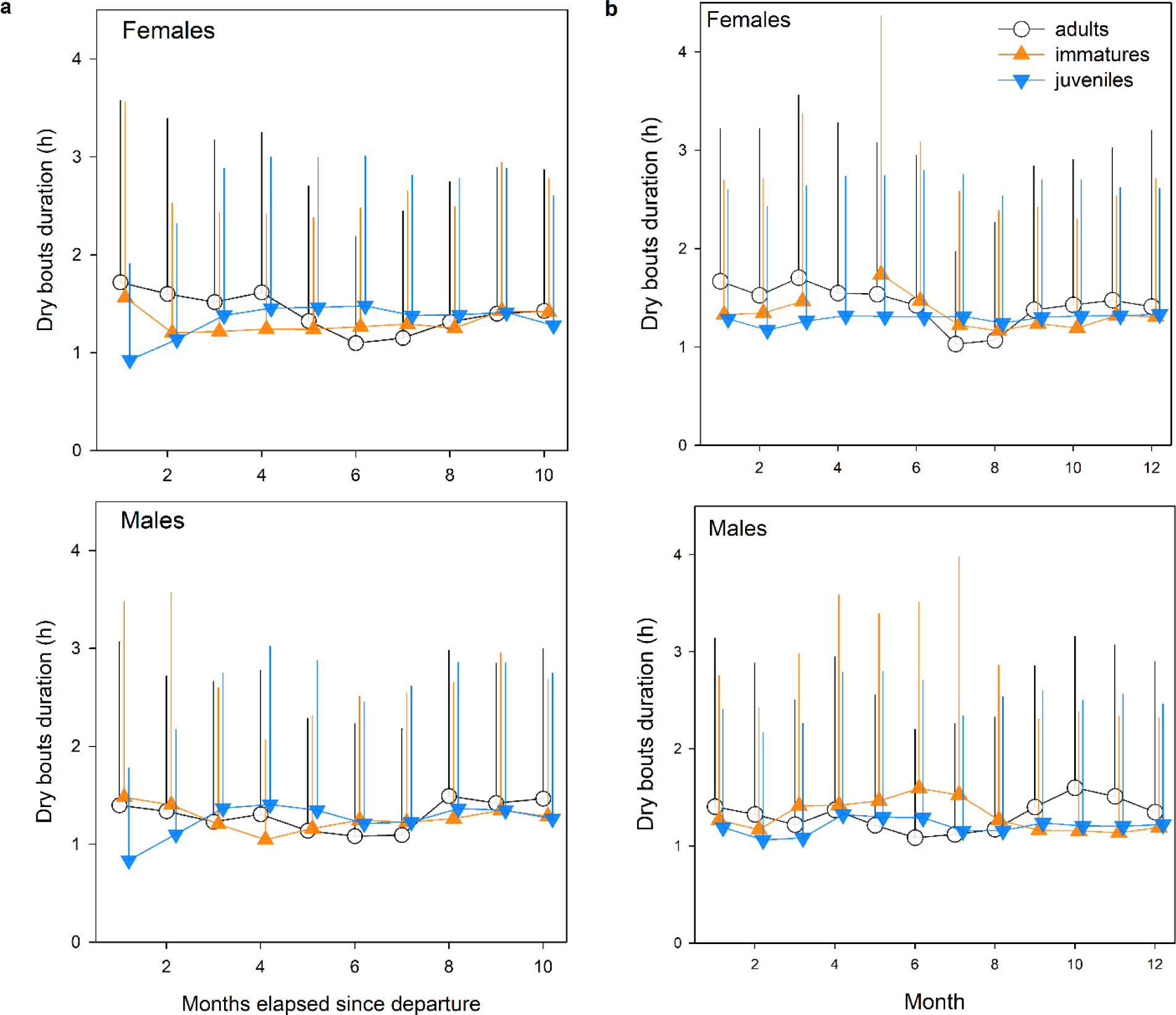
Daily flying bouts duration (dry bouts in hours) depending and on sex (females and males) and on stage (juveniles, immatures and adults) for a) time elapsed since departure from the colony expressed in month (left panel) and for b) each month of the year (right panel). One side error bars represent ± 1 sd

**Figure S3.**
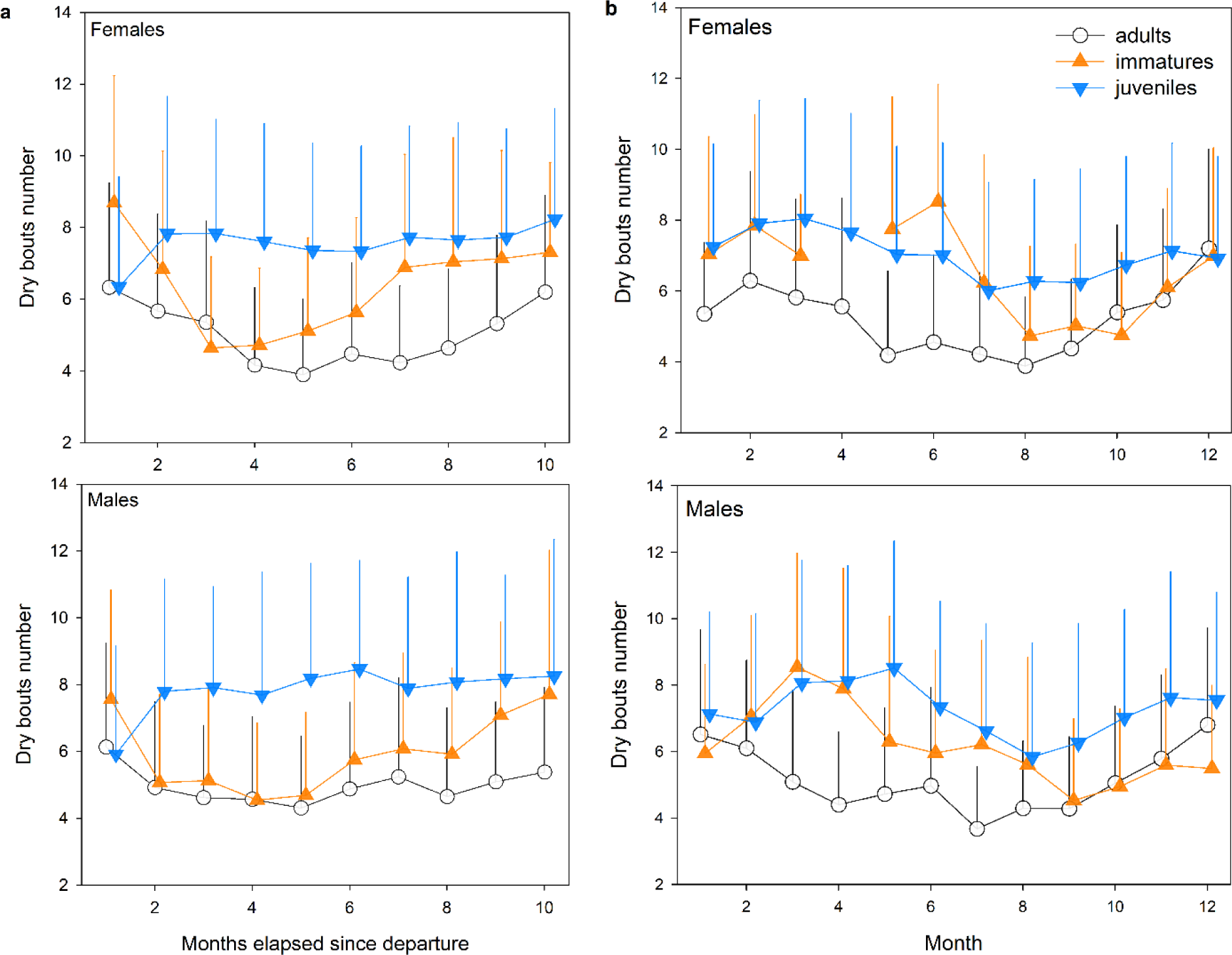
Daily flying bouts number (dry bouts) for every month since departure from the colony for juveniles, immatures and adults for females (upper panel) and males (lower panel). Error bars represent ± 1 sd

**Figure S4.**
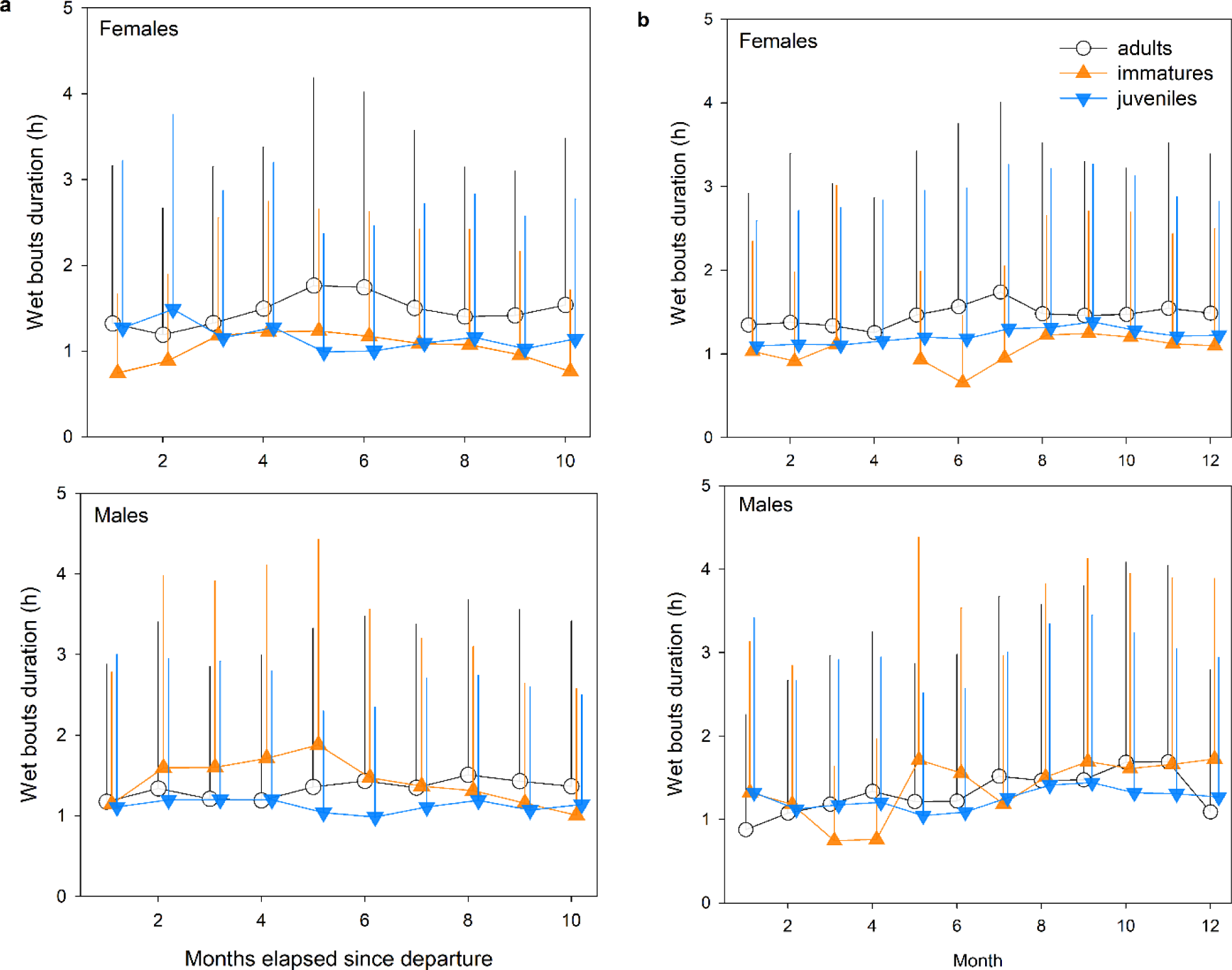
Daily wet bouts duration (bouts on water in hours) depending on stage (juveniles, immatures and adults) and on sex (females and males) for every month since departure of the colony (upper panel) and for each month of the year (lower panel). Error bars represent ± 1 sd

**Figure S5.**
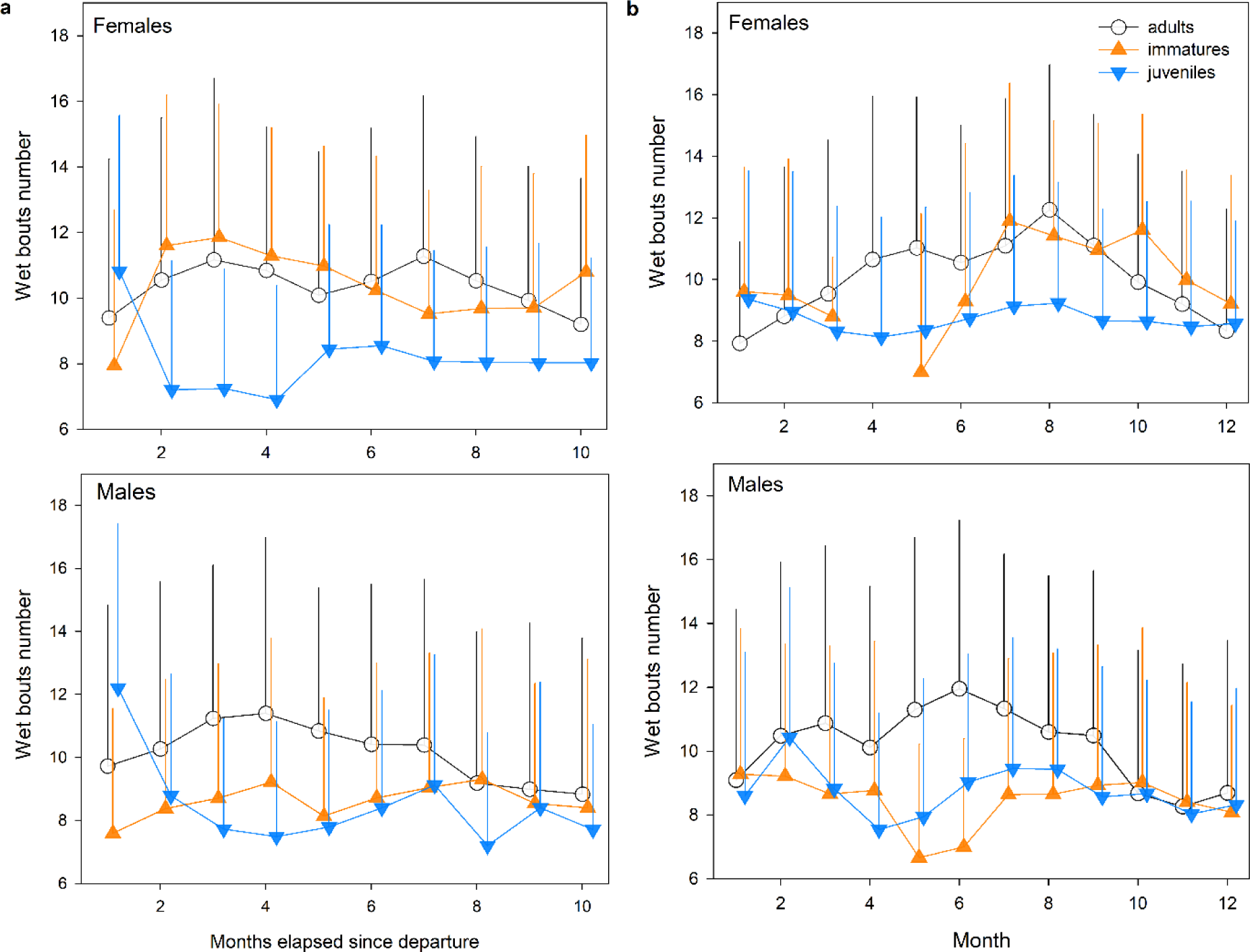
Daily wet bouts number (bouts on water) for every month since departure from the colony for juveniles, immatures and adults for females (upper panel) and males (lower panel). Error bars represent ± 1 sd

**Figure S6.**
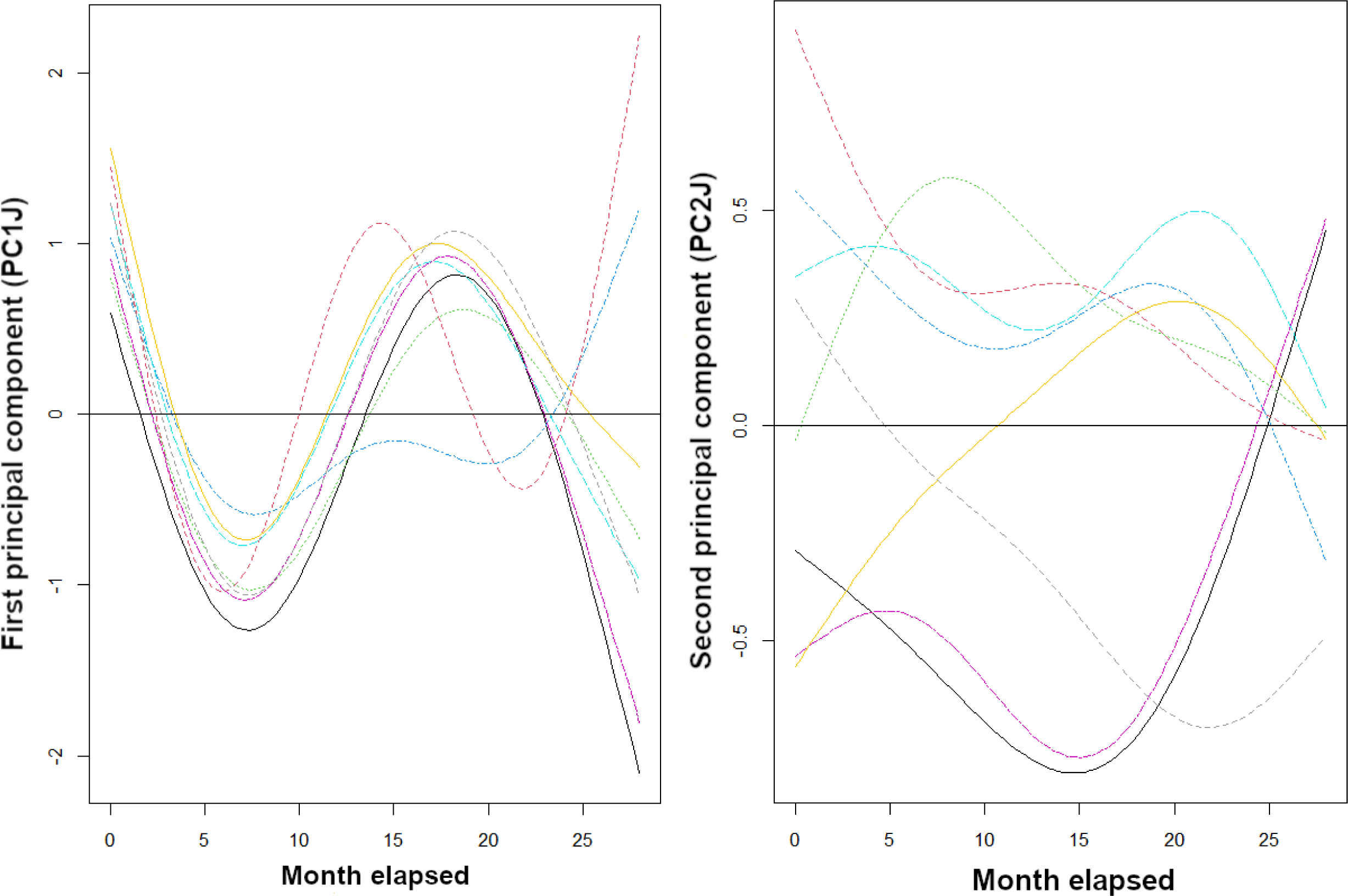
Modeled first (left panel) and second (right panel) axis of principal components analysis of activity parameters of juveniles of Amsterdam albatrosses according to time elapsed (e.g. duration elapsed since departure from the colony expressed in month). Models outputs obtained using random intercepts and slopes (each coloured line representing an individual). Line corresponds to estimated smoother from the GAMM models

**Figure S7.**
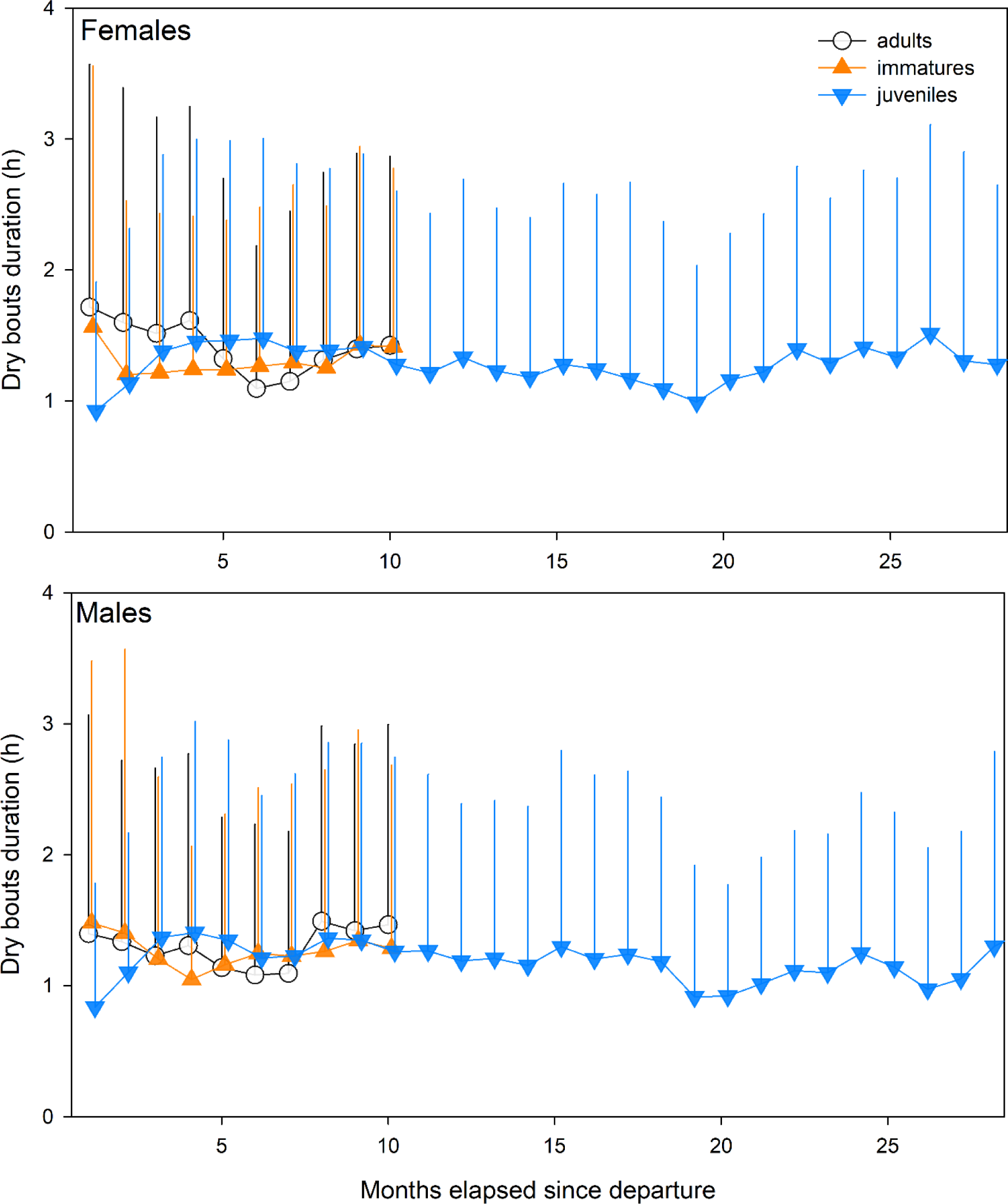
Daily flying bouts duration (dry bouts in hours) for every month since departure of the colony for juveniles, immatures and adults for females (upper panel) and males (lower panel). Error bars represent ± 1 sd

**Figure S8.**
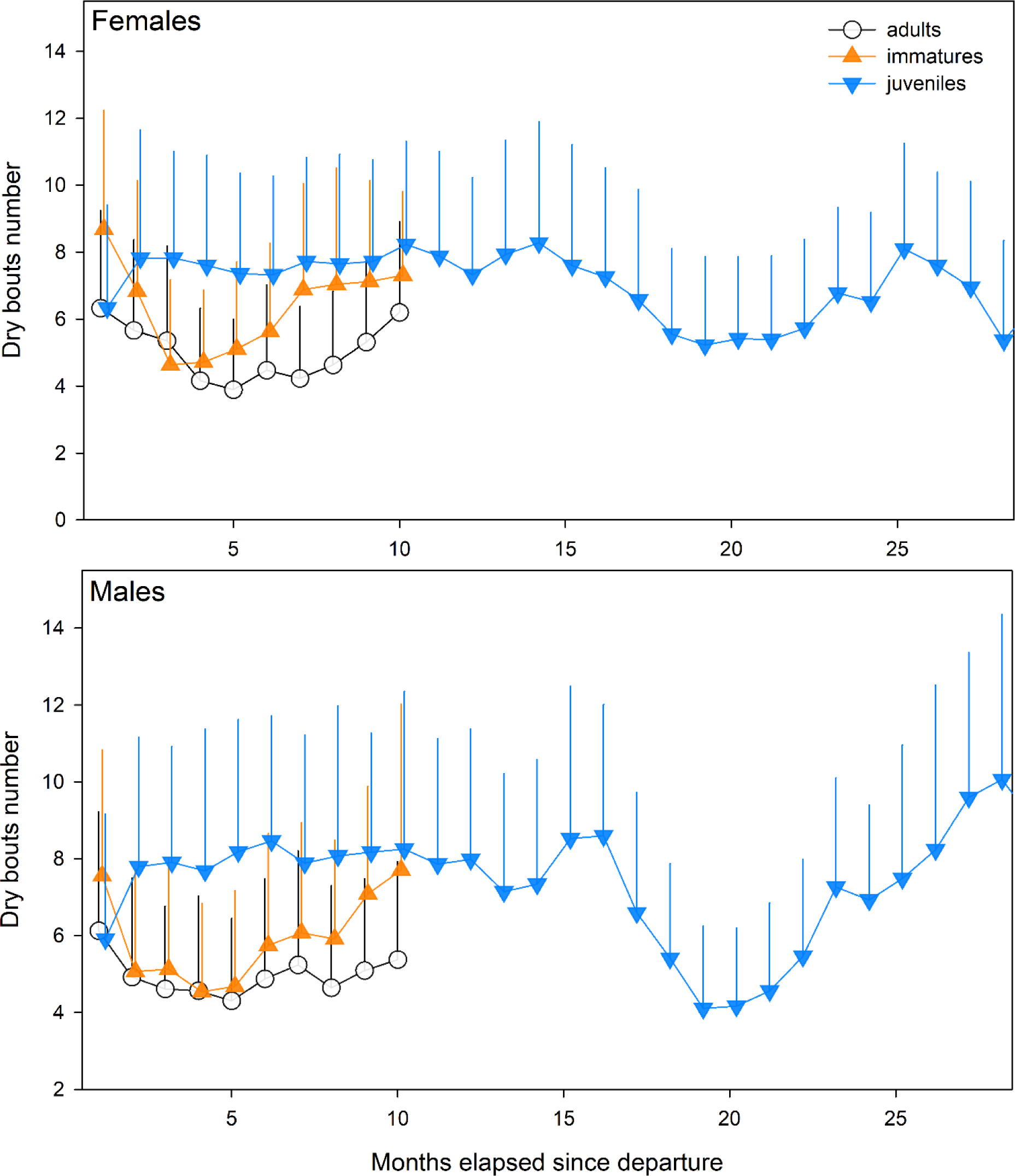
Daily flying bouts number (dry bouts) for every month since departure of the colony for juveniles, immatures and adults for females (upper panel) and males (lower panel). Error bars represent ± 1 sd

**Figure S9.**
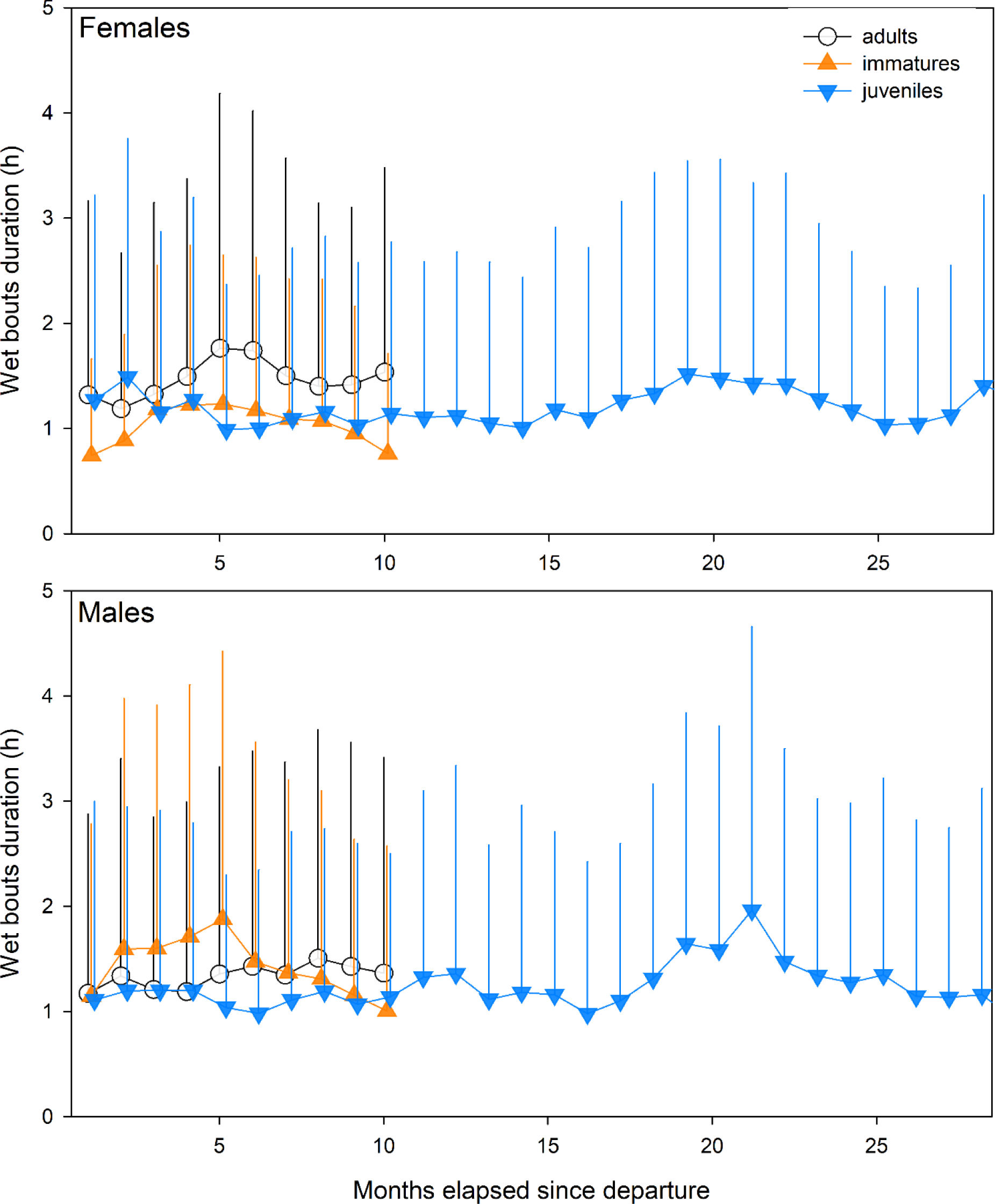
Daily wet bouts duration (bouts on water in hours) for every month since departure of the colony for juveniles, immatures and adults for females (upper panel) and males (lower panel). Error bars represent ± 1 sd

**Figure S10.**
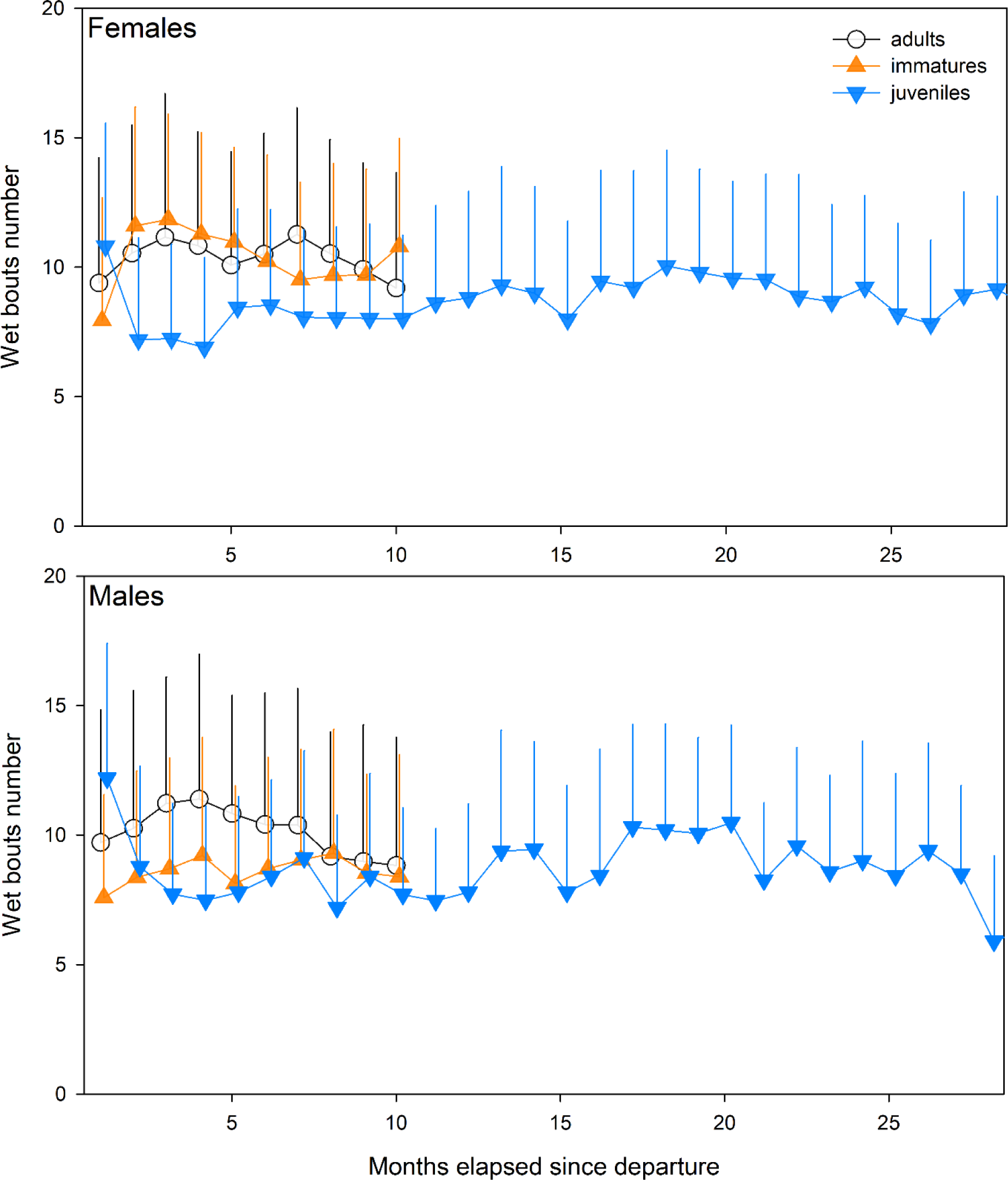
Daily wet bouts number (bouts on water) for every month since departure of the colony for juveniles, immatures and adults for females (upper panel) and males (lower panel). Error bars represent ± 1 sd

**Figure S11.**
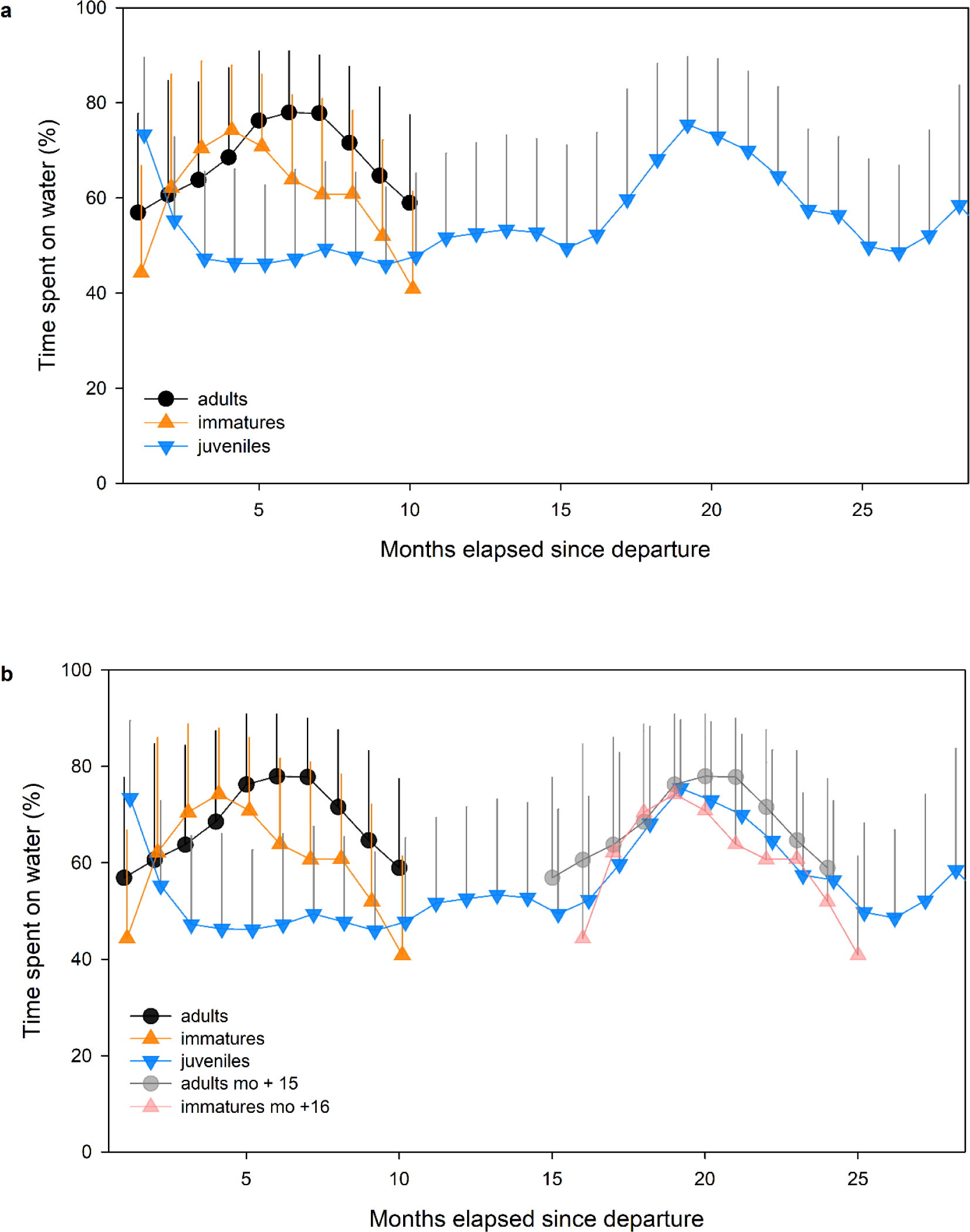
Daily proportions of time spent on water for every month since departure of the colony for juveniles-during the first 28 months spent at sea (after departure), immatures and adults (upper panel) and with a 15-16 months of delay for immatures and adults compared to juveniles (lower panel). Error bars represent ± 1 sd

