## Supplementary material for "The challenges of independence: ontogeny of at-sea behaviour in a long-lived seabird": data-script: ama_activity_pca_R1.r

```
#######################################
###   Script GLM  activite  		###
###   PCA-Multivariate analysis		###
#######################################


# update 24/05/2022
# Delord et al. The challenges of independence: ontogeny of at-sea behaviour in a long-lived seabird 
# R 4.1.3

# 
library("FactoMineR")
library("factoextra")
library("corrplot")
library("Hmisc")

setwd("~/_act")


# dataset for each variable all stage combined
repos<-"~/_act/"
out<-"~/_act/_analyse/_revision/_pca/" 

#1/STADE: toutes donnees regroupees par regroupe_files.r
all.act<- NULL
all.act<-read.csv(file=paste(repos,"ALL-act-data_amal_clean.csv",sep=""),h=TRUE,sep=";")

all.act$stade<-as.factor(all.act$stade)
all.act$device_code<-as.factor(all.act$device_code)
all.act$monthf<-as.factor(all.act$monthf)
all.act$monthelapf<-as.factor(all.act$monthelapf)

active<-all.act[1:8094, 12:16]
head(active)

test<-all.act[1:8094, 1:11]
head(test)

# Correlation matrix
mc_data<-all.act[,2:length(all.act)]
res2<-rcorr(as.matrix(mc_data))
res2
res<-cor(mc_data)
corrplot(res, type = "upper", order = "hclust", tl.col = "black", tl.srt = 45)

# PCA
res.pca<-PCA(active, graph = FALSE)
print(res.pca)
#dimdesc(res.pca)
eig.val<-get_eigenvalue(res.pca)
eig.val
round(res.pca$var$cos2,2)
#dimdesc(res.pca)

png(paste(out,"pca_ama_screenplot.png", sep=""), width=3000, height=1800, res=300)
fviz_eig(res.pca, addlabels = TRUE, ylim = c(0, 50))
dev.off()

var<-get_pca_var(res.pca)
var
head(var$coord, 8)
head(var$cos2, 8)
head(var$contrib, 8)

png(paste(out,"pca_ama_variablesplot.png", sep=""), width=3000, height=1800, res=300)
fviz_pca_var(res.pca, col.var = "black")
dev.off()

png(paste(out,"pca_ama_variablesplot_dim1-2.png", sep=""), width=3000, height=1800, res=300)
fviz_pca_var(res.pca, col.var = "contrib", 
   gradient.cols = c("white", "blue", "red"),
   ggtheme = theme_minimal())
dev.off()
png(paste(out,"pca_ama_variablesplot_dim2-3.png", sep=""), width=3000, height=1800, res=300)
fviz_pca_var(res.pca, axes=c(2,3),col.var = "contrib",
 gradient.cols = c("white", "blue", "red"),
   ggtheme = theme_minimal())
 dev.off()

head(var$cos2, 8)

png(paste(out,"pca_ama_correlation.png", sep=""), width=3000, height=1800, res=300)
corrplot(var$cos2, is.corr = FALSE)
dev.off()

png(paste(out,"pca_ama_cos2-var1-2.png", sep=""), width=3000, height=1800, res=300)
fviz_cos2(res.pca, choice = "var", axes = 1:2)
dev.off()

png(paste(out,"pca_ama_variables-pca.png", sep=""), width=3000, height=1800, res=300)
fviz_pca_var(res.pca, col.var = "cos2",
             gradient.cols = c("#00AFBB", "#E7B800", "#FC4E07"), 
             repel = TRUE # Avoid text overlapping
)
dev.off()

fviz_pca_var(res.pca, alpha.var = "cos2")
head(var$contrib, 8)

png(paste(out,"pca_ama_correlation-dim.png", sep=""), width=3000, height=1800, res=300)
corrplot(var$contrib, is.corr=FALSE)
dev.off()

png(paste(out,"pca_ama_contribution-dim1.png", sep=""), width=3000, height=1800, res=300)
fviz_contrib(res.pca, choice = "var", axes = 1, top = 10)
dev.off()
png(paste(out,"pca_ama_contribution-dim2.png", sep=""), width=3000, height=1800, res=300)
fviz_contrib(res.pca, choice = "var", axes = 2, top = 10)
dev.off()

ind<-get_pca_ind(res.pca)
ind
head(ind$coord, 8)
head(ind$cos2, 8)
head(ind$contrib, 8)

png(paste(out,"pca_ama_individuals.png", sep=""), width=3000, height=1800, res=300)
fviz_pca_ind(res.pca)  
dev.off()          

ind$coord
write.infile(res.pca,file=paste(out,"my_FactoMineR_results.csv",sep=""))   

# 
final<-NULL
final<-cbind(test,ind$coord)

write.csv(final, file=paste(out,"data-pca_amal.csv", sep=""), quote=TRUE, sep=";", row.names=TRUE)     

#-----------------------------------
# 
		rownames(all.act)<-all.act$ID 
		res.pca<-PCA(active, graph = FALSE)

		df_res.pca<- as.data.frame(res.pca$ind$coord) 
		df_res.pca$ID<-rownames(df_res.pca) 

		final<-merge(test, df_res.pca, by="ID" ) 
		write.csv(final, file=paste(out,"data-pca_amal_2.csv", sep=""), quote=TRUE, sep=";", row.names=TRUE)     

#############----------------------------------------------------------------------------
act.group<- NULL
act.group<-read.csv(file=paste(repos,"ALL-act-data_amal_clean_group.csv",sep=""),h=TRUE,sep=";") 

res<-MFA(active2, group=c(1,2,4))

# PCA
res.pca.2<-PCA(all.act, graph = FALSE)

library(cluster)
library(ggplot2)
library(ggfortify)
autoplot(clara(iris[-5], 3))

# PCA
res<-PCA(all.act, quanti.sup= 8:11, quali.sup= 1:7, graph = FALSE)
print(res)
summary(res,nbelements=Inf,file==paste(repos,"essai.txt",sep=""))
plot(res,cex=0.8, shadow=TRUE,habillage=13)

###------------------------------------------------------------------------
#2/JUVENILE: 29 Mo; r
juv.act<- NULL
juv.act<-read.csv(file=paste(repos,"ALL-act-data_juv_amal_clean.csv",sep=""),h=TRUE,sep=";")

juv.act$stade<-as.factor(juv.act$stade)
juv.act$device_code<-as.factor(juv.act$device_code)
juv.act$monthf<-as.factor(juv.act$monthf)
juv.act$monthelapf<-as.factor(juv.act$monthelapf)

active2<-juv.act[1:6161, 12:16]
head(active2)

test2<-juv.act[1:6161, 1:11]
head(test2)

# Correlation matrix
mc_data<-juv.act[,2:length(juv.act)]
res2<-rcorr(as.matrix(mc_data))
res2
res<-cor(mc_data)
corrplot(res, type = "upper", order = "hclust", tl.col = "black", tl.srt = 45)

# PCA
res.pca<-PCA(active2, graph = FALSE)
print(res.pca)
#dimdesc(res.pca)
eig.val<-get_eigenvalue(res.pca)
eig.val
round(res.pca$var$cos2,2)
#dimdesc(res.pca)

png(paste(out,"pca_ama_juv_screenplot.png", sep=""), width=3000, height=1800, res=300)
fviz_eig(res.pca, addlabels = TRUE, ylim = c(0, 50))
dev.off()

var<-get_pca_var(res.pca)
var
head(var$coord, 8)
head(var$cos2, 8)
head(var$contrib, 8)

png(paste(out,"pca_ama_juv_variablesplot.png", sep=""), width=3000, height=1800, res=300)
fviz_pca_var(res.pca, col.var = "black")
dev.off()

png(paste(out,"pca_ama_juv_variablesplot_dim1-2.png", sep=""), width=3000, height=1800, res=300)
fviz_pca_var(res.pca, col.var = "contrib", 
   gradient.cols = c("white", "blue", "red"),
   ggtheme = theme_minimal())
dev.off()
png(paste(out,"pca_ama_juv_variablesplot_dim2-3.png", sep=""), width=3000, height=1800, res=300)
fviz_pca_var(res.pca, axes=c(2,3),col.var = "contrib",
 gradient.cols = c("white", "blue", "red"),
   ggtheme = theme_minimal())
 dev.off()

head(var$cos2, 8)

png(paste(out,"pca_ama_juv_correlation.png", sep=""), width=3000, height=1800, res=300)
corrplot(var$cos2, is.corr = FALSE)
dev.off()

png(paste(out,"pca_ama_juv_cos2-var1-2.png", sep=""), width=3000, height=1800, res=300)
fviz_cos2(res.pca, choice = "var", axes = 1:2)
dev.off()

png(paste(out,"pca_ama_juv_variables-pca.png", sep=""), width=3000, height=1800, res=300)
fviz_pca_var(res.pca, col.var = "cos2",
             gradient.cols = c("#00AFBB", "#E7B800", "#FC4E07"), 
             repel = TRUE 
)
dev.off()

fviz_pca_var(res.pca, alpha.var = "cos2")
head(var$contrib, 8)

png(paste(out,"pca_ama_juv_correlation-dim.png", sep=""), width=3000, height=1800, res=300)
corrplot(var$contrib, is.corr=FALSE)
dev.off()

png(paste(out,"pca_ama_juv_contribution-dim1.png", sep=""), width=3000, height=1800, res=300)
fviz_contrib(res.pca, choice = "var", axes = 1, top = 10)
dev.off()
png(paste(out,"pca_ama_juv_contribution-dim2.png", sep=""), width=3000, height=1800, res=300)
fviz_contrib(res.pca, choice = "var", axes = 2, top = 10)
dev.off()

ind<-get_pca_ind(res.pca)
ind
head(ind$coord, 8)
head(ind$cos2, 8)
head(ind$contrib, 8)

png(paste(out,"pca_ama_juv_individuals.png", sep=""), width=3000, height=1800, res=300)
fviz_pca_ind(res.pca)  
dev.off()          

ind$coord
write.infile(res.pca,file=paste(out,"my_FactoMineR_juv_results.csv",sep=""))   #to export a list

# 
final<-NULL
final<-cbind(test2,ind$coord)

write.csv(final, file=paste(out,"data-pca_juv_amal.csv", sep=""), quote=TRUE, sep=";", row.names=TRUE)     

#-----------------------------------
# 
		rownames(all.act)<-all.act$ID 
		res.pca<-PCA(active, graph = FALSE)

		df_res.pca<- as.data.frame(res.pca$ind$coord) 
		df_res.pca$ID<-rownames(df_res.pca) 

		final<-merge(test, df_res.pca, by="ID" ) 
		write.csv(final, file=paste(out,"data-pca_amal_2.csv", sep=""), quote=TRUE, sep=";", row.names=TRUE)     
```
