## Supplementary material for "The challenges of independence: ontogeny of at-sea behaviour in a long-lived seabird": data-script: ama_revision_act_gamm_juveniles_R1.r

```
###############################
###   Script GAMM  activite  ###
###   mixed model structure; random effect on id
###############################

# update 27/5/22 revision article PCI 
# input data = PCA axes values
# Delord et al. The challenges of independence: ontogeny of at-sea behaviour in a long-lived seabird 

library("lme4")
library("nlme")
library("stats")
require("lubridate")
require("ggplot2")
### GAMM voir gamm_itsadug.docx
library("gamm4")
library("MuMIn")
library(itsadug)
# explo analysis
# testing different models
library(mgcv)
library(MASS)


## ------------------------------------------------------------------------
setwd("~/_act/_revision")
gls.out<-"~/_revision/"

## ------------------------------------------------------------------------
# data PCA JUVENILES
## ------------------------------------------------------------------------
#
input <- NULL
input<-read.table("~/_revision/_pca/data-pca_juv_amal_modif.csv",sep=";",h=T) 
# format
input$jour<-as.POSIXct(strptime(input$jour,format="%d/%m/%Y", "GMT"))
input$stade<-as.factor(input$stade)
input$device_code<-as.factor(input$device_code)
input$monthf<-as.factor(input$monthf)
input$sex<-as.factor(input$sex)

df2<-input

# A.1/ AXE 1: dim1 juveniles
df2 <- na.omit(df2) 
 
Cj.0<-gam(Dim.1 ~ 1+s(monthelap,device_code, bs='re'), data=df2, family = gaussian)
summary(Cj.0) 
print(Cj.0)
AIC((Cj.0))
 
Cj.1<-gam(Dim.1 ~ s(monthelap,k=2)+s(monthelap,device_code, bs='re'), data=df2, family = gaussian)
summary(Cj.1)
print(Cj.1)
AIC((Cj.1))

Cj.2<-gam(Dim.1 ~ s(monthelap,k=2)+monthf+s(monthelap,device_code, bs='re'), data=df2, family = gaussian) ## BEST
summary(Cj.2)
print(Cj.2)
AIC((Cj.2))
gam.check(Cj.2)
gam.vcomp(Cj.2)

Cj.3<-gam(Dim.1 ~ s(monthelap,k=2)+monthf+sex+s(monthelap,device_code, bs='re'), data=df2, family = gaussian)
summary(Cj.3)
print(Cj.3)
AIC((Cj.3))

par(mfrow=c(2,2)) ; gam.check(Cj.2)
par(mfrow=c(1,1)); plot(Cj.2)

##___________________________________________________________________
		b<-gam(Dim.1 ~ s(monthelap, k = 2),  na.action="na.omit", data=df2)
		plot(b,pages=1,residuals=TRUE) ## hint of a problem in s(x2)
		## the following suggests a problem with s(x2)
		summary(b)
		gam.check(b)
		k.check(b, subsample=5000, n.rep=400)
		rsd <- residuals(b)
		gam(rsd~s(monthelap,k=2,bs="cs"),gamma=1.4,data=df2) ## fine
		gam(rsd~s(monthelap,k=6,bs="cs"),gamma=1.4,data=df2) ## fine
		
##___________________________________________________________________
		library(mgcViz) # https://cran.r-project.org/web/packages/mgcViz/vignettes/mgcviz.html
		b <- getViz(Cj.2) #extract the first smooth component using the sm function
				o <- plot( sm(b, 1) )
				Cj.2.monthelap.fit<-o$data$fit	

out<-("~/_act/_revision/")
write.csv(Cj.2.monthelap.fit, "ama_pca-gammDim1_juveniles_monthelap-fitCj2_revised.csv")

# A.2/ AXE 2: dim2 juveniles
df2 <- na.omit(df2) 
 
Cj2.0<-gam(Dim.2 ~ 1, data=df2, family = gaussian)
Cj2.0<-gam(Dim.2 ~ 1+s(monthelap,device_code, bs='re'), data=df2, family = gaussian)
summary(Cj2.0) 
print(Cj2.0)
AIC((Cj2.0))

Cj2.1<-gam(Dim.2~ s(monthelap,k=2)+s(monthelap,device_code, bs='re'),
            family=gaussian(),data=df2)
summary(Cj2.1)
print(Cj2.1)
AIC((Cj2.1))

Cj2.2<-gam(Dim.2 ~ s(monthelap,k=2)+monthf+s(monthelap,device_code, bs='re'), data=df2, family = gaussian) 
summary(Cj2.2)
print(Cj2.2)
AIC((Cj2.2))
gam.check(Cj2.2)

Cj2.3<-gam(Dim.2 ~ s(monthelap,k=2)+monthf+sex+s(monthelap,device_code, bs='re'), data=df2, family = gaussian) ## BEST
summary(Cj2.3)
print(Cj2.3)
AIC((Cj2.3)) # BEST

par(mfrow=c(2,2)) ; gam.check(Cj2.3)

par(mfrow=c(1,1)); plot(Cj2.3)

		#library(mgcViz) # https://cran.r-project.org/web/packages/mgcViz/vignettes/mgcviz.html
		b <- getViz(Cj2.3) #extract the first smooth component using the sm function
		# month elapsed
				o <- plot( sm(b, 1) )
				Cj2.3.monthelap.fit<-o$data$fit	

write.csv(Cj2.3.monthelap.fit, "ama_pca-gammDim2_juveniles_monthelap-fitCj2.3_revised.csv")

# A.3/ AXE 3: dim3 juveniles
df2 <- na.omit(df2) 
 
Cj3.0<-gam(Dim.3 ~ 1, data=df2, family = gaussian)
summary(Cj3.0) 
print(Cj3.0)
AIC((Cj3.0))

Cj3.0r<-gam(Dim.3 ~ 1+s(monthelap,device_code, bs='re'), data=df2, family = gaussian)
summary(Cj3.0r) 
AIC((Cj3.0r))

Cj3.1<-gam(Dim.3~ s(monthelap,k=2)+s(monthelap,device_code, bs='re'),
            family=gaussian(),data=df2)
summary(Cj3.1)
print(Cj3.1)
AIC((Cj3.1))

Cj3.2<-gam(Dim.3 ~ s(monthelap,k=2)+monthf+s(monthelap,device_code, bs='re'),
            family=gaussian(),data=df2)
summary(Cj3.2)
print(Cj3.2)
AIC((Cj3.2))
gam.check(Cj3.2)

Cj3.3<-gam(Dim.3 ~ monthf+s(monthelap,device_code, bs='re'),family=gaussian(),data=df2)
summary(Cj3.3)
print(Cj3.3)
AIC((Cj3.3))

Cj3.4<-gam(Dim.3 ~ monthf+sex+s(monthelap,device_code, bs='re'), data=df2, family = gaussian) ## BEST
summary(Cj3.4)
print(Cj3.4)
AIC((Cj3.4))

Cj3.5<-gam(Dim.3 ~ s(monthelap,k=2,bs="cs")+monthf+sex,random = ~(1+ monthelap|device_code),
            family=gaussian(),data=df2,method="REML")
summary(Cj3.5)
print(Cj3.5)
AIC((Cj3.5))

par(mfrow=c(1,1)); plot(Cj3.4)

		#library(mgcViz) # https://cran.r-project.org/web/packages/mgcViz/vignettes/mgcviz.html
		b <- getViz(Cj3.4) #extract the first smooth component using the sm function
		# month elapsed
				o <- plot( sm(b, 1) )
				Cj3.4.monthelap.fit<-o$data$fit	

write.csv(Cj3.4.monthelap.fit, "ama_pca-gammDim3_juveniles_monthelap-fitCj3.4_revised.csv")

# 

m4 <- bam(Dim.3 ~ s(monthelap,k=2,bs="cs") + s(monthelap, device_code, bs='fs', k=5), data=df2, discrete=TRUE)
summary(m4)
par(mfrow=c(1,2)) ; plot(m4)

par(mfrow=c(1,2), cex=1.1)
plot(m4, select=2)
abline(h=0)
inspect_random(m4, select=2) #Inspection and interpretation of random factor smooths. 
```
