## Supplementary material for "The challenges of independence: ontogeny of at-sea behaviour in a long-lived seabird": data-script: ama_revision_act_gamm_stades_R2.r

```
#######################################################
###   Script GAMM  activite  						###
###   mixed model structure; random effect on id	###
#######################################################

# update 15/5/23 revision #2 article PCI Ecology
# input data = PCA axes values
# Delord et al. The challenges of independence: ontogeny of at-sea behaviour in a long-lived seabird 

library("lme4")
library("nlme")
library("stats")
require("lubridate")
require("ggplot2")
library("gamm4")
library("MuMIn")
library(itsadug)
# explo analysis
# testing different models
library(mgcv)
library(MASS)


## ------------------------------------------------------------------------
setwd("~/_revision")
gls.out<-"~/_revision/"

## ------------------------------------------------------------------------
# data PCA STADES
input <- NULL
input<-read.table("~/_revision/_pca/data-pca_amal.csv",sep=";",h=T) 
# format
input$jour<-as.POSIXct(strptime(input$jour,format="%d/%m/%Y", "GMT"))
input$stade<-as.factor(input$stade)
input$device_code<-as.factor(input$device_code)
input$monthf<-as.factor(input$monthf)
input$sex<-as.factor(input$sex)

l1<-input

# A.1/ AXE 1: dim1 all stages
#
library(fitdistrplus)
dim1<-l1$Dim.1	#	descdist(dim1, discrete = FALSE)

l1 <- na.omit(l1) 

# MODEL 
		 # model null
		 Cm.0<-gam(Dim.1 ~ 1, data=l1, family = gaussian)
		summary(Cm.0)
		print(Cm.0)
		AIC((Cm.0))

		# model null + random
		 Cm.0r<-gam(Dim.1 ~ 1+s(monthelap,device_code, bs='re'), data=l1, family = gaussian)
		summary(Cm.0r) 
		print(Cm.0r)
		AIC((Cm.0r))
		 
		Cm.1<-gam(Dim.1 ~ s(monthelap,k=2)+s(monthelap,device_code, bs='re'),
					family=gaussian(),data=l1,method="REML")
		summary(Cm.1)
		print(Cm.1)
		AIC(Cm.1)

		Cm.2<-gam(Dim.1 ~ s(monthelap,k=2)+monthf+s(monthelap,device_code, bs='re'),
					family=gaussian(),data=l1,method="REML")
		summary(Cm.2)
		print(Cm.2)
		AIC(Cm.2)

		Cm.3<-gam(Dim.1 ~ s(monthelap,k=2)+monthf+stade+s(monthelap,device_code, bs='re'),
					family=gaussian(),data=l1,method="REML")
		summary(Cm.3)
		print(Cm.3)
		AIC((Cm.3))

		Cm.4<-gam(Dim.1 ~ s(monthelap,k=2)+monthf+stade+sex+s(monthelap,device_code, bs='re'),
					family=gaussian(),data=l1,method="REML")
		summary(Cm.4)
		print(Cm.4)
		AIC((Cm.4)) 

		par(mfrow=c(1,1)); plot(Cm.4)

		Cm.5<-gam(Dim.1 ~ s(monthelap, by=stade, bs="cs")+monthf+stade+sex,random = ~(1+ monthelap|device_code),
					family=gaussian(),data=l1,method="REML")
		summary(Cm.5)
		print(Cm.5)
		AIC((Cm.5)) 
		par(mfrow=c(2,2)) ; gam.check(Cm.5)

##___________________________________________________________________
		b<-gam(Dim.1 ~ s(monthelap, k = 2),  na.action="na.omit", data=l1)
		plot(b,pages=1,residuals=TRUE) ## hint of a problem in s(x2)
		## the following suggests a problem with s(x2)
		summary(b)
		gam.check(b)
		k.check(b, subsample=5000, n.rep=400)
		rsd <- residuals(b)
		gam(rsd~s(monthelap,k=2,bs="cs"),gamma=1.4,data=l1) ## fine
		gam(rsd~s(monthelap,k=6,bs="cs"),gamma=1.4,data=l1) ## fine
##___________________________________________________________________
		library(mgcViz) # https://cran.r-project.org/web/packages/mgcViz/vignettes/mgcviz.html
		b <- getViz(Cm.4) #extract the first smooth component using the sm function
		# month elapsed
				o <- plot( sm(b, 1) )
				Cm.4.monthelap.fit<-o$data$fit	

out<-("~/_act/_revision/")
write.csv(Cm.4.monthelap.fit, file=paste(out,"ama_pca-gammDim1_Moulting_monthelap-fit_revised.csv",sep=""))

# A.2/ AXE 2: dim2 all stages
l1 <- na.omit(l1) 
 
# MODEL  
		 # model null
		 Cm2.0<-gam(Dim.2 ~ 1, data=l1, family = gaussian)
		summary(Cm2.0) 
		print(Cm2.0)
		AIC((Cm2.0))

		# model null + random
		 Cm2.0r<-gam(Dim.2 ~ 1+s(monthelap,device_code, bs='re'), data=l1, family = gaussian)
		summary(Cm2.0r) 
		print(Cm2.0r)
		AIC((Cm2.0r))

		Cm2.1<-gam(Dim.2 ~ s(monthelap,k=3)+s(monthelap,device_code, bs='re'),
					family=gaussian(),data=l1,method="REML")
		summary(Cm2.1)
		print(Cm2.1)
		AIC((Cm2.1))

		Cm2.2<-gam(Dim.2 ~ s(monthelap,k=3)+monthf+s(monthelap,device_code, bs='re'),
					family=gaussian(),data=l1,method="REML")
		summary(Cm2.2)
		print(Cm2.2)
		AIC((Cm2.2))

		Cm2.3<-gam(Dim.2 ~ s(monthelap,k=3)+monthf+stade+s(monthelap,device_code, bs='re'),
					family=gaussian(),data=l1,method="REML") # BEST
		summary(Cm2.3)
		print(Cm2.3)
		AIC((Cm2.3))

		Cm2.4<-gam(Dim.2 ~ s(monthelap,k=3)+monthf+stade+sex+s(monthelap,device_code, bs='re'),
					family=gaussian(),data=l1,method="REML")
		summary(Cm2.4)
		print(Cm2.4)
		AIC((Cm2.4)) 

		par(mfrow=c(2,2)) ; gam.check(Cm2.3)
		par(mfrow=c(1,1)); plot(Cm2.3)

		Cm2.5<-gam(Dim.2 ~ s(monthelap,by=stade)+monthf+stade,random = ~(1+ monthelap|device_code),
					family=gaussian(),data=l1,method="REML")
		summary(Cm2.5)
		print(Cm2.5)
		AIC((Cm2.5))

##___________________________________________________________________
		b<-gam(Dim.2 ~ s(monthelap, k = 4),  na.action="na.omit", data=l1)
		plot(b,pages=1,residuals=TRUE) ## hint of a problem in s(x2)
		## the following suggests a problem with s(x2)
		summary(b)
		gam.check(b)
		k.check(b, subsample=5000, n.rep=400)
		rsd <- residuals(b)
		gam(rsd~s(monthelap,k=2,bs="cs"),gamma=1.4,data=l1) ## fine
		gam(rsd~s(monthelap,k=6,bs="cs"),gamma=1.4,data=l1) ## fine
##___________________________________________________________________
		library(mgcViz) # https://cran.r-project.org/web/packages/mgcViz/vignettes/mgcviz.html
		b <- getViz(Cm2.3) #extract the first smooth component using the sm function
				o <- plot( sm(b, 1) )
				Cm2.3.monthelap.fit<-o$data$fit	

write.csv(Cm2.3.monthelap.fit, "ama_pca-gammDim2_stades_monthelap-fitCm2.3_update.csv")

# A.3/ AXE 3: dim3 all stages
l1 <- na.omit(l1) 
 
# MODEL 
		# model null
		 Cm3.0<-gam(Dim.3 ~ 1, data=l1, family = gaussian)
		summary( Cm3.0) #ne plotter que les relations signif et changer les y
		print( Cm3.0)
		AIC(( Cm3.0))

		# model null + random
		 Cm3.0r<-gam(Dim.3 ~ 1+s(monthelap,device_code, bs='re'), data=l1, family = gaussian)
		summary(Cm3.0r) #ne plotter que les relations signif et changer les y
		print(Cm3.0r)
		AIC((Cm3.0r))

		Cm3.1<-gam(Dim.3 ~ s(monthelap,k=2)+s(monthelap,device_code, bs='re'),
					family=gaussian(),data=l1,method="REML")
		summary(Cm3.1)
		print(Cm3.1)
		AIC((Cm3.1))

		Cm3.2<-gam(Dim.3 ~ monthf+s(monthelap,device_code, bs='re'),
					family=gaussian(),data=l1,method="REML")
		summary(Cm3.2)
		print(Cm3.2)
		AIC((Cm3.2))

		Cm3.3<-gam(Dim.3 ~ monthf+stade+s(monthelap,device_code, bs='re'),
					family=gaussian(),data=l1,method="REML")
		summary(Cm3.3)
		print(Cm3.3)
		AIC((Cm3.3))

		Cm3.4<-gam(Dim.3 ~ monthf+stade+sex+s(monthelap,device_code, bs='re'),
					family=gaussian(),data=l1,method="REML")# BEST
		summary(Cm3.4)
		print(Cm3.4)
		AIC((Cm3.4))

		par(mfrow=c(2,2)) ; gam.check(Cm3.4)

		par(mfrow=c(1,1)); plot(Cm3.4)

		Cm3.5<-gam(Dim.3 ~ s(monthelap,k=3)+monthf+stade+sex+s(monthelap,device_code, bs='re'),
					family=gaussian(),data=l1,method="REML")
		summary(Cm3.5)
		print(Cm3.5)
		AIC((Cm3.5)) 
		par(mfrow=c(1,1)); plot(Cm3.5)

		Cm3.6<-gam(Dim.3 ~ s(monthelap,by=stade)+monthf+stade+sex,random = ~(1+ monthelap|device_code),
					family=gaussian(),data=l1,method="REML")
		summary(Cm2.5)
		print(Cm2.5)
		AIC((Cm2.5))

##___________________________________________________________________
		b<-gam(Dim.3 ~ s(monthelap, k = 4),  na.action="na.omit", data=l1)
		plot(b,pages=1,residuals=TRUE) ## hint of a problem in s(x2)
		## the following suggests a problem with s(x2)
		summary(b)
		gam.check(b)
		k.check(b, subsample=5000, n.rep=400)
		rsd <- residuals(b)
		gam(rsd~s(monthelap,k=1,bs="cs"),gamma=1.4,data=l1) ## fine
		gam(rsd~s(monthelap,k=3,bs="cs"),gamma=1.4,data=l1) ## fine
		
##___________________________________________________________________
		library(mgcViz) # https://cran.r-project.org/web/packages/mgcViz/vignettes/mgcviz.html
		b <- getViz(Cm3.4) #extract the first smooth component using the sm function
				o <- plot( sm(b, 1) )
				Cm3.4.monthelap.fit<-o$data$fit	
write.csv(Cm3.4.monthelap.fit, "ama_pca-gammDim3_stades_monthelap-fitCm3.4_update.csv")
```
